# Tissue specificity and chromosomal alterations shape divergent immune programs in HRD tumors

**DOI:** 10.1101/2025.10.13.681986

**Authors:** Doga C. Gulhan, David Barras, Marco Mina, Eleonora Ghisoni, Yoo-Na Kim, Vinay Viswanadham, Hu Jin, Florian Huber, Krisztian Homicsko, Michal Bassani-Sternberg, Giovanni Ciriello, Peter J. Park, Denarda Dangaj Laniti

## Abstract

Homologous recombination deficiency (HRD) activates pro-inflammatory cGAS/STING signaling, positioning it as a biomarker for combining immune checkpoint blockade (ICB) and PARP inhibition (PARPi). However, the consequences of HRD on the immune landscape across cancers remain unclear. Here, we applied a pan-cancer HRD classifier to >10,000 tumors from The Cancer Genome Atlas and uncovered striking heterogeneity in immune activity. Compared to HR-proficient tumors, HRD tumors showed elevated inflammation in breast, ovarian, and endometrial cancers. These tumors exhibited robust activation of innate and adaptive immune pathways (IFN, NF-κB) and transcriptional hallmarks of senescence, angiogenesis, and adenosine signaling. In contrast, lung, head and neck, and melanoma HRD tumors displayed suppressed inflammation and evidence of immune escape through large-scale loss-of-heterozygosity (LOH) at IFNA/B, STING, and other loci. These tumors also frequently presented HLA LOH and oncogene amplifications, suggesting selection under immune pressure and replication stress. Together, our study resolves HRD tumors into two immune archetypes, immune-inflamed and immune-evasive, linked to chromosomal instability and lineage, informing biomarker-driven evaluation of immune checkpoint blockade/PARPi combinatorial therapies.

## Introduction

DNA repair deficiencies have been associated with enhanced anti-tumor immune responses and are therefore considered key biomarkers for immune checkpoint blockade (ICB) response. For example, mismatch repair deficiency (MMRD) increases the generation of neoantigens^1,2^ and activates cytosolic DNA sensing through the cGAS/STING pathway^3,4^, leading to robust type I interferon production and T cell infiltration. These features underlie the high immunogenicity and ICB responsiveness of MMRD tumors^5^. Similarly, homologous recombination deficiency (HRD), often caused by *BRCA1* or *BRCA2* loss, can promote tumor-intrinsic inflammation via STING and interferon (IFN) signaling and upregulate chemokines such as *CCL5* and *CXCL9*, thereby enhancing immunogenicity and T cell inflammation^6–10^. Consequently, HRD has emerged as a promising biomarker for predicting response to ICB therapy^11^, particularly when combined with PARP inhibition^12,13^. While numerous studies in breast and ovarian cancers support an association between HRD and heightened immunogenicity^7,9,14,15^, it remains unclear whether this relationship generalizes across cancer types. To determine whether HRD universally promotes immune activation or instead drives divergent immune programs across tumor types, a systematic pan-cancer evaluation is required.

A comprehensive pan-cancer analysis of the immune landscape in HRD requires assembling a large, well-defined cohort of HRD tumors. Although biallelic *BRCA1*/*2* loss is the canonical indicator, many HRD tumors lack these alterations^16^, motivating complementary detection methods. Current classification strategies therefore combine (1) pathogenic mutations or promoter hypermethylation in additional DNA double-strand break (DSB) repair genes^17^, and (2) genomic “scar” measures such as loss of heterozygosity (LOH)^18^, telomeric allelic imbalance^19^, and large-scale state transitions^20^, as summarized in the genomic instability score (GIS) by the *MyChoice CDx*^21^ HRD assay (Myriad Genetics). More recently, mutational signature analysis^22,23^ has revealed that HRD tumors bear distinctive single-base substitution (SBS) patterns, most notably signature 3 (Sig3). We previously developed *SigMA*, a computational tool that detects the HRD-associated single-base substitution signature (Sig3) even from low-mutation-count datasets such as targeted gene panels^12,31,32^, enabling improved prediction of PARP inhibitor (PARPi) response even in *BRCA1*/*2*-wildtype tumors^12,32^. In addition to point mutations, HRD tumors bear characteristic indel and rearrangement features (>5 bp deletions at microhomologies; rearrangement signatures with 1-10 kb deletions and duplications^24–28^). Because these signatures can be robustly extracted from whole-genome sequencing (WGS) data, WGS-based HRD classification can be performed with high confidence^29,30^; however, current WGS pan-cancer cohorts are still limited in size, and the lack of transcriptional profiling in most WGS tumors restricts the investigation of immune characteristics of HRD tumors. Therefore, our first step below was to develop a new HRD classification approach for whole-exome data, which enabled us to analyze a large cohort.

Through integrated genomic, transcriptomic, and immune deconvolution analyses, we ask three related questions. First, is HRD linked to a single, uniform immune state or to distinct, lineage- and context-dependent programs across cancers? Second, which genomic determinants appear to modulate immune activity in HRD tumors, and how do pervasive chromosomal instability features such as loss of heterozygosity (LOH) or copy-number amplification affect their immune properties? Third, how does HRD relate to the tumor microenvironment (TME), including the composition of myeloid and lymphoid populations, and to transcriptional programs implicated in inflammation, senescence, angiogenesis, and adenosine signaling that shape the TME?

To address these questions, we derived a composite inflammation score (IS) from bulk RNA-seq that captures cGAS–STING–associated type-I interferon responses, chemokine-driven T-cell recruitment, IFN-γ signaling, and cytotoxic CD8⁺ T-cell activation. We orthogonally validate this score in an independent cohort with matched bulk RNA-seq and immunofluorescence, then apply it across TCGA to compare HRD and HR proficient (HRP) tumors within each cancer type. In parallel, we perform cell-state deconvolution of the TME from bulk RNA-seq using single-cell–derived reference signatures to quantify immune and stromal populations. We integrate copy-number/LOH landscapes with immune readouts to test whether gene-dosage at immune-regulatory loci or oncogenes tracks with inflammatory state. Through this integrative approach, we define immune archetypes of HRD tumors, pinpoint their defining properties, and assess their prevalence across cancer types.

Together, our findings define a comprehensive map of immune features associated with HRD across cancers and reveal how lineage context and chromosomal instability sculpt the immunologic consequences of HRD, providing a foundation for biomarker-guided combination strategies that rationally pair PARP inhibition with immunotherapy.

## Results

### Pan-cancer HRD classification

Although pan-cancer HRD classification methods based on WGS have been developed^29,33^, a similar approach tailored for whole-exome sequencing (WES) data was needed to dramatically increase the number of samples we can analyze. Because the number of variants detected in WES is an order of magnitude smaller than in WGS, a more robust approach is needed. Therefore, we developed a machine learning method named *CSI-HRD* (<u>C</u>NA, <u>S</u>BS, and Indel-based <u>HRD</u> Classifier) (**Figure 1A, Figure S1A**). This method employs gradient-boosting classifiers trained both on pan-cancer data and individually by tumor type, leveraging the increased sample size of pan-cancer cohorts while accommodating tissue-specific differences across cancer types. As one of the input features, we utilized Sig3 scores from the latest version of the *SigMA* algorithm,^31^ optimized for TCGA datasets^34^. We combined the *SigMA* score with GIS (estimated from paired SNP array data) and counts of small deletions of at least 5bp occurring with and without overlap at microhomologies. These two indel counts reflect the indel signatures ID6 and ID8, linked with microhomology-mediated and non-homologous end-joining double-strand break repair processes, respectively, and shown to be enriched in HRD tumors^29^ (**Figure S1B**). Based on the *CSI-HRD* scores from the pan-cancer and tumor type-specific classifiers, we labeled tumors with high scores for both classifiers as HRD and those with low scores for both classifiers as HRP; the remaining 3.5% of samples in between were categorized as HRD-low (**Figure S1C, Methods**). MMRD and POLE proofreading-deficient samples were grouped into separate categories (**Figure 1B and C**). To validate our HRD classification, we compared our WES-based classification to HRD status calculated from WGS data^35^ (using the HRDetect algorithm^29^) for a subset of samples for which both WES and WGS data were available, showing 91.5% concordance (**Figure S1D, E;** 797 samples with published WGS-based HRDetect calls^35^). We also confirmed that the majority of *BRCA1/2*^−/−^ samples were in the HRD category (**Figure S1F**). HRD-low or HRP *BRCA1/2*^−/−^ samples (59 out of 1066; 5.5% of all HRD-labeled cases) were labeled HRD independently of their *CSI-HRD* classification status. The final pan-cancer labels for the entire TCGA dataset are provided in **Table S1**, as a resource for future studies.

**Figure 1.**
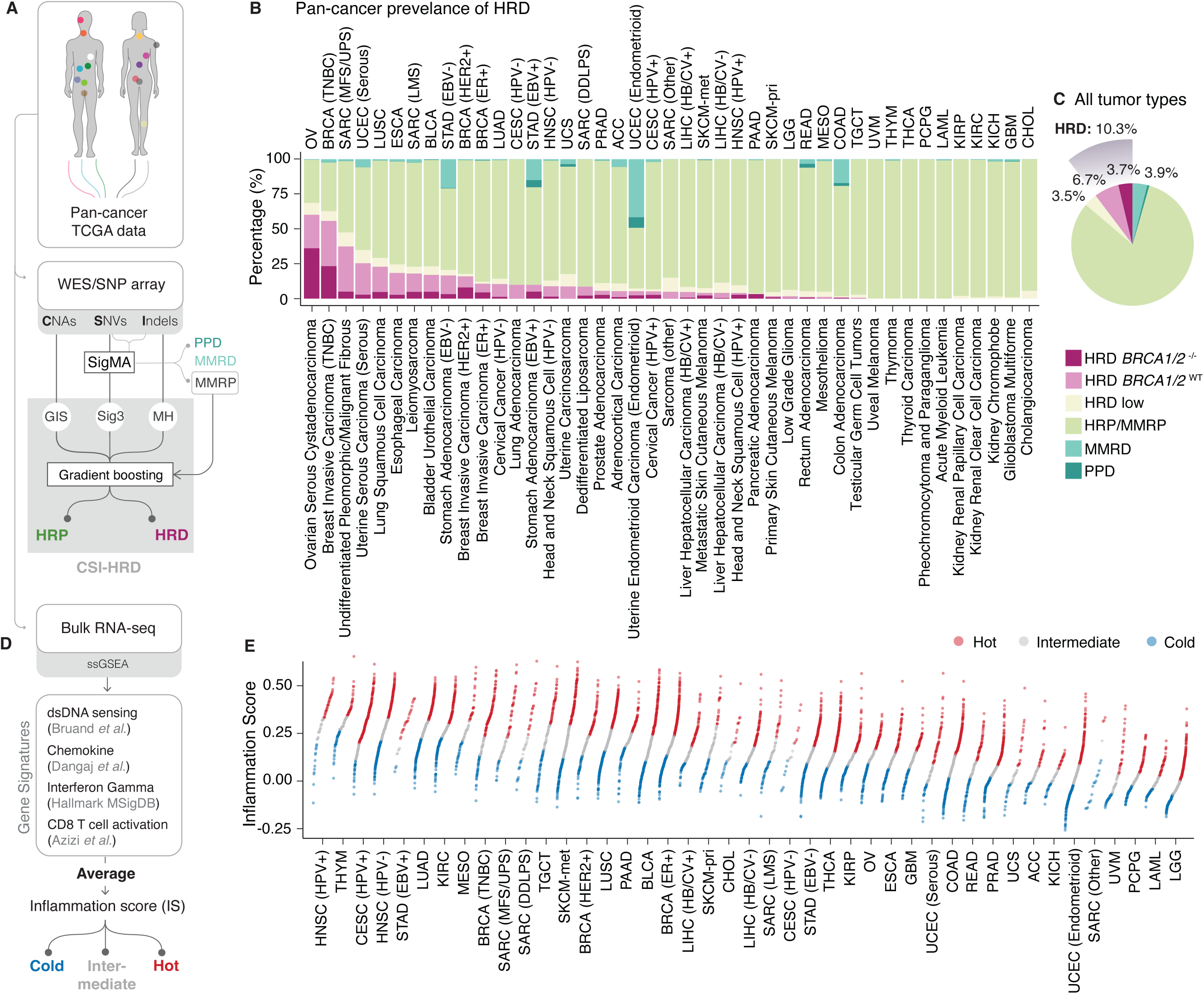
Pan–Cancer Characterization of HRD and Inflammation. **(A)** Pan–cancer HRD classification workflow. **(B)** Percentage of samples in HRD (split into *BRCA1*/*2*^⁻/⁻^ and *BRCA1*/*2*^WT^), HRP/MMRP, MMRD, and PPD categories, sorted by descending HRD prevalence. Cancer type abbreviations appear at the top and full names at the bottom. **(C)** Prevalence of the same categories as in panel B across pan–cancer data. **(D)** Inflammation score (IS), defined as the average of ssGSEA scores for four gene sets; samples are stratified into hot, intermediate, and cold categories based on IS tertiles within each tumor type. Tumors with viral infection, MMRD, or PPD status were excluded when calculating tertiles. **(E)** Distribution of IS in each tumor type. Each point displays a sample colored according to IS category (hot, intermediate, and cold). Tumor types are ordered in descending order for mean IS.

As anticipated, the prevalence of HRD differed across tumor types (**Figure 1B**), with an overall incidence of 10.5% in pan-cancer data. Ovarian (OV) and triple-negative breast cancers (TNBCs) had the highest rates of HRD (59.3% and 55.8%, respectively). Strikingly, 64.6% of HRD tumors in the pan-cancer dataset lacked BRCA1/2 mutations (*BRCA1*/2^WT^), underscoring the significance of our HRD classification method (**Figure 1C**). *BRCA1*/*2*^−/−^ cases accounted for approximately half (50.6%) of HRD cases in tissues recognized for germline *BRCA1*/*2*-linked cancer susceptibility (e.g. OV: Ovarian serous cystadenocarcinoma, TNBC, ER^+^ or HER2^+^ BRCA: Breast invasive carcinoma samples, PAAD: Pancreatic adenocarcinoma, PRAD: Prostate adenocarcinoma). In contrast, *BRCA1*/*2*^−/−^ tumors represented a substantially smaller fraction (21.9%) of HRD cases in other cancer types.

To uncover additional mechanisms that contribute to HRD beyond *BRCA1/2* loss, we examined whether other DNA damage response (DDR) genes (**Table S2;** compiled using Reactome^36^ and Ref. ^37^) were selectively altered in HRD compared to HRP samples, apart from the *BRCA1*/*2*^−/−^ cases. We included genetic alterations (including SBSs and indels predicted to be truncating, pathogenic, or damaging, as well as deep deletions) and epigenetic silencing through promoter hypermethylation in these comparisons. We observed that 83 out of 410 examined DDR genes were significantly enriched in *BRCA1*/*2*^WT^ HRD compared to HRP samples (**Table S2**), including *BRIP1*, *BLM, FANCF*, *FANCM, PALB2*, and *RAD51C* (**Figure S2A)**. To explore potential epistatic interactions, we performed enrichment analyses focused on gene pairs involving DDR genes or *TP53*, testing whether HRD was enriched specifically when both genes were dysfunctional. Through this analysis, we identified 14 additional DDR genes significantly enriched in HRD, consistently paired with TP53 (**Table S3**). The only exception was *HERC2*, which demonstrated significant co-occurrence with *ATR* and *PRKDC* as well as *TP53*. These paired alterations were significantly enriched in *CSI-HRD*-classified tumors (Fisher’s exact test; *p* < 0.0001) (**Figure S2B**). Notably, approximately 25% of HRD-classified tumors lacked detectable alterations in DDR genes (**Figure S2C**), yet exhibited hallmark features of HRD, elevated Sig3 scores, high GIS, and increased frequency of microhomology-mediated small deletions (**Figure S2D**).

Our findings highlight the strength of our mutational signature–based classifiers *CSI-HRD* for detecting HRD across diverse cancer types, even in the absence of canonical *BRCA1/2* alterations. By integrating multiple genomic features from WES and SNP array data, *CSI-HRD* offers a scalable and clinically relevant tool for pan-cancer HRD detection without requiring WGS, enabling more inclusive patient stratification for HRD-targeted therapies.

### HR deficiency shapes anti-tumor proinflammatory signaling in a cancer type– specific manner

Building on this refined pan-cancer HRD classification, we next investigated whether HRD status is associated with pro-inflammatory signaling within the tumor microenvironment (TME), and whether such immunogenic effects are omnipresent in all cancer types. We developed an inflammation score (IS) based on bulk RNA-seq data designed to capture cGAS/STING-mediated pro-inflammatory signaling and T cell infiltration and activation (**Figure 1D and E**). The IS score (**Table S1**) was calculated as the average of single-sample gene set enrichment analysis (ssGSEA) scores using four curated gene signatures reflecting: (i) type-I IFN response triggered by cytosolic DNA sensing, (ii) intratumoral recruitment of T cells via release of chemokines, (iii) IFN-γ response, and (iv) cytotoxic CD8^+^ T cell activation^8,9,38,39^.

To validate our IS, we applied it to an independent cohort of immune-classified treatment-naïve ovarian cancers for which we had previously generated both bulk RNA-seq and high-resolution whole-slide immunofluorescence (IF) imaging of CD8^+^ tumor-infiltrating lymphocytes (TIL)^10^. Our IS robustly captured both CD8^+^ TIL abundance and their spatial distribution within the tumor epithelium; thus, distinguishing inflamed tumors with high CD8^+^ TIL levels within the tumor epithelium from excluded ones where CD8^+^ T cells are primarily observed in the stroma (**Figure S3A, B**). Based on the IS distribution within each tumor type, we defined ‘hot’ (more inflamed) and ‘cold’ (less inflamed) categories, using the upper and lower tertiles of the IS within each tumor type of the TCGA dataset, respectively (**Figure 1E**).

Next, we compared IS between HRD and HRP tumors within each cancer type and revealed striking heterogeneity in the association between HR deficiency and tumor inflammation across cancers. HRD tumors exhibited significantly higher IS than their HRP counterparts in ER^+^ BRCA, OV, and endometrioid uterine corpus endometrial carcinomas (UCEC-Endometrioid) (**Figure 2A**). We termed these tumor types deficient-Hot/proficient-Cold (dHpC), as the observed pattern may result from either heightened inflammation in HRD tumors or reduced inflammation in HRP tumors. The tumor types with a non-significant (n.s.) increase in IS in HRD compared to HRP were referred to as dHpC-n.s. In contrast, HRD tumors displayed lower inflammation than HRP tumors in human papillomavirus-negative (HPV^−^) head-neck squamous cell carcinoma (HNSC), lung adenocarcinoma (LUAD), lung squamous cell carcinoma (LUSC), and metastatic skin cutaneous melanoma (SKCM-met). We classified those as deficient-Cold/proficient-Hot (dCpH) cancer types (**Figure 2A**). To validate these distinct inflammatory patterns, we compared IS values of archetypal HRD tumors harboring *BRCA1/2^−/^*^−^ alterations with HRP tumors; the differences were significant in both groups (**Figure 2A**; Z-score transformed IS comparison in dHpC *p* = 0.021, t = 2.3, in dCpH *p* = 0.027, t = −2.3; t-test), confirming that the interplay between HRD and inflammation is highly tumor-type specific.

**Figure 2.**
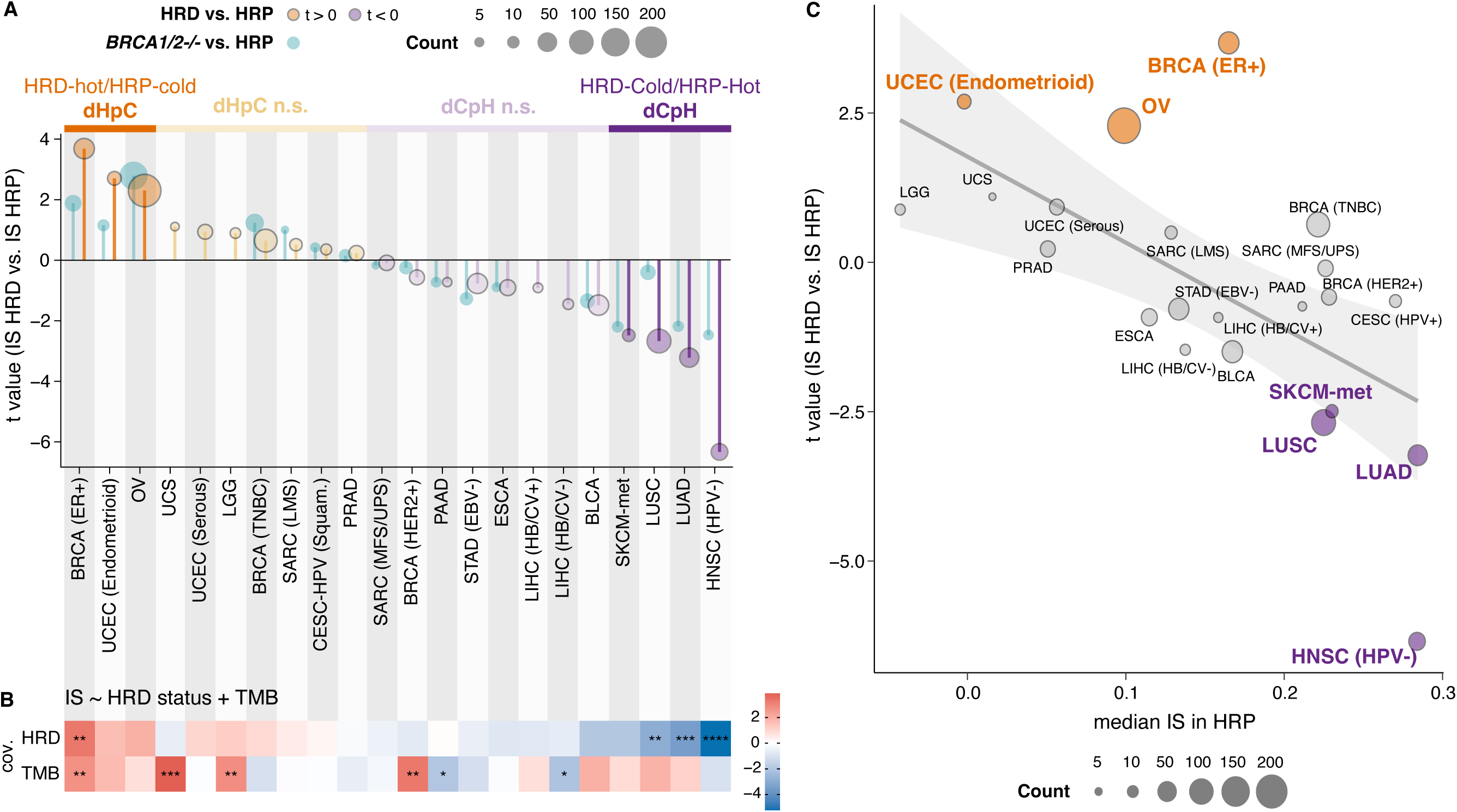
Differential inflammation between HRD and HRP tumors across pan–cancer data. **(A)** T values comparing IS between HRD and HRP/MMRP tumors (orange for t > 0; purple for t < 0), with darker shades indicating p < 0.05. t values for *BRCA1*/*2*^⁻/⁻^ versus HRP/MMRP tumors are shown in cyan. Tumor-type groups are labeled dHpC (deficient hot/proficient cold) or dCpH (deficient cold/proficient hot) when HRD tumors exhibit higher or lower IS, respectively; the suffix “n.s.” denotes non-significant differences. Point size reflects the number of HRD samples. **(B)** T values from a multiple linear regression of IS on HRD status and tumor mutational burden (TMB). Asterisks denote p values: *p < 0.05; **p < 0.01; ***p < 0.001; ****p < 0.0001. **(C)** Scatter plot of the t values from panel A versus median IS in HRP samples (baseline IS per tissue). Fill colors denote dHpC and dCpH groups; a linear regression line with 95% confidence intervals (gray shading) is shown.

To uncover factors contributing to the tissue-specific relationship between HRD and inflammation, we examined TMB^40–43^ as a potential confounder. Notably, tumor types in the dCpH group (melanoma, lung, and head and neck cancers) are characterized by high TMB, unlike those in the dHpC group (**Figure S3C**). When comparing TMB in HRD and HRP categories within each tumor type group, we found that TMB was higher in HRD cancers (**Figure S3D**). This led us to conclude that TMB differences cannot explain the lower IS observed in the dCpH cancer group. To further disentangle the relationship, we performed multilinear regression analyses incorporating both TMB and HRD status as covariates. These models confirmed that HRD was independently associated with lower IS in dCpH tumors and higher IS in dHpC tumors (**Figure 2B, Figure S3E**), re-affirming that differences in TMB do not explain the tumor-type specific nature of the HRD-inflammation axis.

The dCpH group – comprising tumor types where HRD tumors exhibit lower inflammation than their HRP counterparts – includes cancers with inherently high immunogenicity, such as lung cancer and metastatic melanoma. This pattern suggests that in these tumors, HRD may promote immune evasion rather than immune activation. We hypothesized that increased and selective immune pressure on HRD tumors in highly immunogenic microenvironments could underlie the observed heterogeneity across tumor types. Supporting this hypothesis, we observed an inverse correlation between the median IS of HRP tumors and the t-values from HRD vs HRP comparisons across all tumor types (**Figure 2C**, *p* = 0.0017; r = −0.63; Pearson’s correlation).

These findings reveal that the immunogenic consequences of HR deficiency are not universal but shaped by the immune context of each cancer type. In immune-hot tumors such as melanoma and lung cancer, HRD may be counter-selected through immune pressure, promoting immune evasion. In contrast, in immune-cold tumors such as ovarian or ER^+^ breast cancers, HRD is linked to enhanced inflammation, supporting its potential as a predictive biomarker for combinatorial immunotherapy.

### Divergent immune microenvironments in HRD tumors reveal coordinated and suppressive TME archetypes across cancer types

Given the pronounced tumor-type-specific heterogeneity in the relationship between HR deficiency and inflammation, we sought to systematically characterize the immune and stromal components of the tumor microenvironment (TME) in HRD and HRP tumors. We aimed to identify tumor-extrinsic factors that may modulate the inflammatory landscape associated with HRD. To achieve this, we performed immune and stromal cell deconvolution from bulk RNA-seq data using CIBERSORTx^44^ with single-cell (sc) RNA-seq-derived immune signatures^45^ (**Table S1; Methods**). This approach enabled high-resolution inference of immune cell populations and states, providing greater specificity than previous bulk-RNAseq-based analyses^46^. We Z-score transformed the relative abundance of each inferred immune cell population within tumor types and calculated the difference between HRD and HRP tumors, as well as between hot and cold-HRD subsets. Hierarchical clustering of immune and stromal cell populations based on the comparisons between HRD and HRP and across hot and cold-HRD tumors resulted in the identification of 5 distinct, but potentially co-occurring groups (**Figure 3A**).

**Figure 3.**
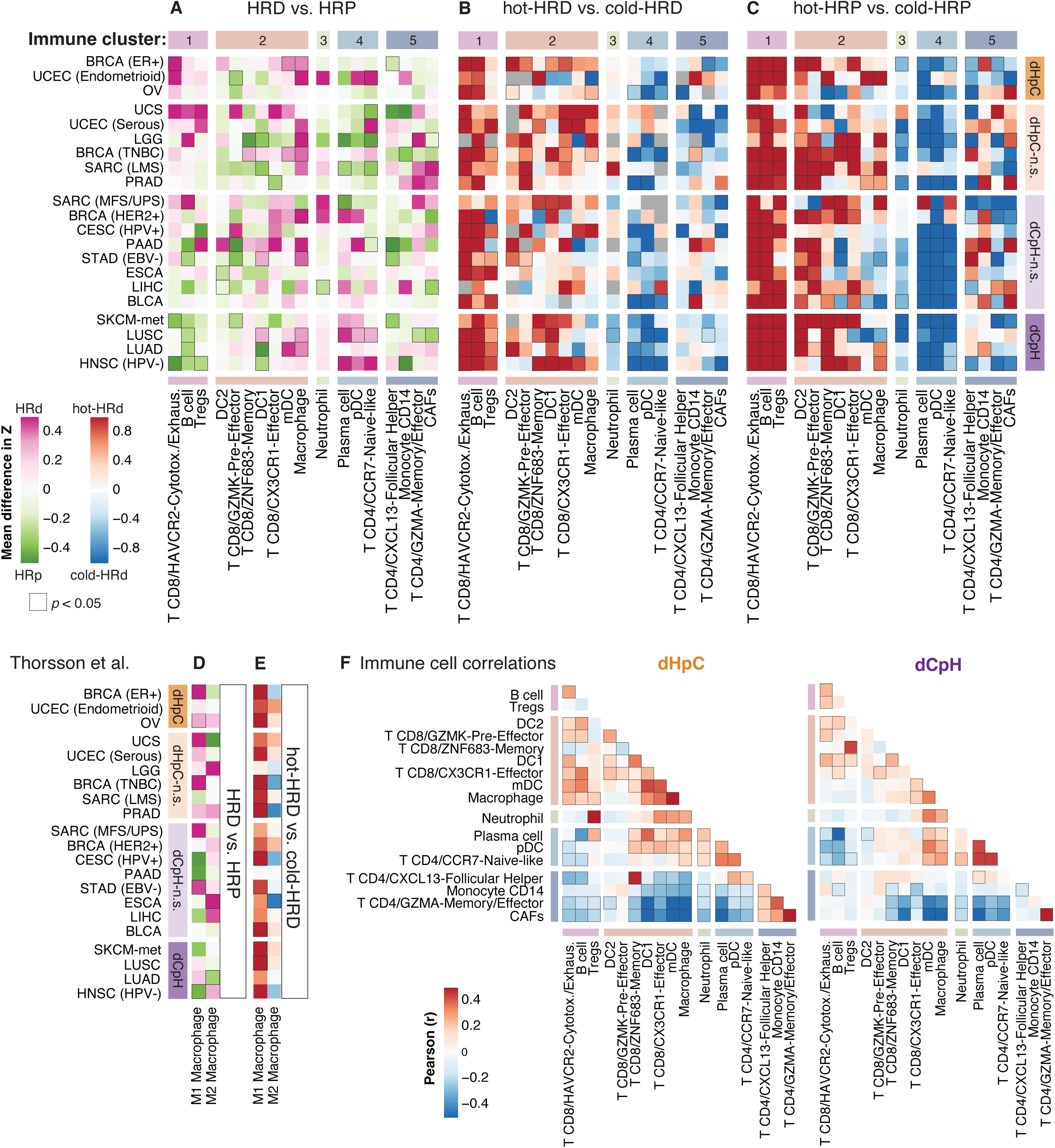
Tumor-Type-Specific TME Archetypes in HRD versus HRP Tumors. **(A**-**C)** Heatmap of differences in relative proportions of CIBERSORTx-inferred immune and stromal cells between HRD and HRP/MMRP (**A**) tumors, between hot HRD and cold HRD tumors (**B**) or between hot HRP and cold HRP tumors (**C**) across cancer types, clustered into five modules (clusters 1–5). The colors show the ΔZ difference between the groups after Z-score normalization. Cluster 1-5 of immune cell types were obtained by columns-based hierarchical clustering of ΔZ values from all three comparisons (**A**-**C**). **(D**-**E)** Heatmap of ΔZ values for M1 and M2 macrophage scores between HRD and HRP tumors (**D**) or between hot HRD and cold HRD tumors (**E**). Tumor types appear in the same order than in Figure 2A (descending t value for IS comparison); significant changes (p < 0.05 by t-test) are marked with black borders. **(F)** Pearson correlation between Z-normalized relative immune cell proportions in dHpC (left) and dCpH (right) groups; *p* < 0.05 are indicated by black borders around tiles.

In dCpH, tumor-reactive TILs in cluster 1 were depleted in HRD tumors compared to their HRP counterparts, whereas the opposite trend was observed in dHpC cancer groups, where cluster 1 cells were enriched in HRD tumors (**Figure 3A; Figure S4A, B**). This cluster included cytotoxic/exhausted HAVCR2^+^ CD8^+^ T cells expressing effector molecules (*GZMB*, *GNLY*, *IFNG*), inhibitory markers (*HAVCR2*, *PDCD1*, *CTLA-4*, *LAG3*, *TIGIT*) as well as *CXCL13*, along with regulatory T cells (Tregs), and B cells. These cell types have established roles in antigen recognition, immune reactivity, and immune regulation^47–49^. Consistent with their immune-reactive identity, cluster 1 cells were universally more abundant in immune-hot compared to immune-cold tumors, irrespective of HRD status, across all examined tumor types (**Figure 3B, C**). Their abundance correlated strongly with IS values, particularly for cytotoxic/exhausted CD8^+^ T cells (Pearson; r = 0.57, *p* < 2.2e-16; **Figure S4C**). The enrichment of exhausted CD8^+^ T cells and Tregs in dHpC likely reflects active tumor recognition by the immune system, followed by exhaustion and immune regulation, making these cancers potential candidates for immune checkpoint blockade. Conversely, their depletion in dCpH HRD tumors may indicate impaired immune recognition, and a more pronounced immune-evasive phenotype, consistent with our previous observations (**Figure 2C**).

Cluster 2 comprised diverse myeloid cell types, including macrophages, dendritic cells (DC1, DC2, and myeloid dendritic cells (mDCs)), as well as GZMK^+^ CD8^+^ pre-effector, CX3CR1^+^ CD8^+^ effector, and ZNF683^+^ CD8^+^ memory T cells, subsets previously linked to immunotherapy responses^50,51^. This cluster represents key T cell-myeloid niches^52^ that may play an essential role in shaping the TME of HRD tumors^10,52^. Across tumor types, macrophages and mDCs from cluster 2 were enriched in HRD compared to HRP tumors (**Figure 3A**). M1 macrophage^46^ scores were elevated with HRD in most dHpC and dHpC-n.s. cancers (**Figure 3D**), but no such enrichment was observed in dCpH and dCpH-n.s. cases (**Figure 3D**). M1 macrophages were also more prevalent in hot-HRD tumors across all tumor types **(Figure 3E**), underscoring a strong association between HRD-driven inflammation and pro-inflammatory innate immune activity. These findings align with previous reports showing that tumor-intrinsic IFN–JAK–STAT signaling, triggered by DNA sensing and STING activation, can polarize innate immune cells and promote inflammatory myeloid responses^9,14,15,53^. Importantly, in dHpC HRD tumors, the relative abundance of mDCs and macrophages correlated with tumor-reactive cell populations from cluster 1, suggesting a coordinated immune TME landscape driven by HRD in these settings (**Figure 3F**). In contrast, this correlation was absent in dCpH tumors (**Figure 3F**), reinforcing the more immune-evasive phenotype of HRD tumors in cancers that are inherently highly immunogenic.

We also identified two clusters, clusters 4 and 5, that were preferentially enriched in cold HRP tumors. Cluster 4 comprised plasmacytoid dendritic cells (pDCs), plasma cells, and naïve-like CD4^+^ T cells, which were more abundant in dCpH HRD tumors compared with their HRP counterparts (**Figure 3A**). These cell types were enriched in cold-HRD tumors, particularly in the dCpH group as well as in cold-HRP tumors (**Figure 3C**). Cluster 4 abundance was also associated with cluster 3 (**Figure 3F**), which was dominated by neutrophils, suggesting coordinated infiltration of non-activated or potentially suppressive immune populations in these tumors.

Cluster 5 consisted of cancer-associated fibroblasts (CAFs), CD14^+^ monocytes, CXCL13^+^CD4^+^ follicular helper cells, and GZMA^+^ CD4^+^ memory/effector T cells (**Figure 3A**). Among these, CAFs were notably enriched in HRD tumors within specific cancer types, such as prostate adenocarcinoma (PRAD) and certain sarcomas (SARC-LMS, and SARC-MFS/UPS), relative to HRP counterparts (**Figure 3A**). This CAF accumulation may promote immune cell exclusion, potentially explaining the lack of significant IS enrichment in HRD versus HRP tumors in these settings (**Figure 2**). Although the precise role of Cluster 4 cells in anti-tumor immunity remains unclear, we observed a predominantly inverse correlation between the abundance of Clusters 4 and 5, suggesting they represent distinct immunosuppressive TME phenotypes (**Figure 3F**).

Together, these results reveal that HRD tumors exhibit diverse and context-dependent immune microenvironments, ranging from inflamed, immune-responsive niches (characterized by CD8^+^ TILs, mDCs, and M1 macrophages in dHpC cancers) to immune-excluded or inert phenotypes (dominated by CAFs, neutrophils, and naïve lymphoid cells) in dCpH and stromal-rich cancers. The presence or absence of coordinated immune remodeling appears tightly linked to tumor histology and immunogenicity, with critical implications for predicting which HRD tumors may respond to immune checkpoint blockade.

### Chromosomal instability drives immune evasion in HRD tumors via disruption of interferon signaling and amplification of oncogenic stress pathways

The reduced immune activation in dCpH prompted us to investigate potential tumor-intrinsic mechanisms of immune evasion in HRD tumors of these histotypes. To identify and quantify these mechanisms, we compared the genomic landscapes of HRD tumors across cancer types.

Previous pan-cancer studies, focusing primarily on SNVs and indels, have identified multiple immune escape mechanisms in immunologically cold tumors, including loss-of-function alterations in antigen presentation genes (e.g., *HLA* and *B2M*), *PTEN* and JAK/STAT alterations, and dysregulation of WNT/b-catenin pathways^46,54–56^. Aneuploidy can influence multiple immune-related genes and has been reported as a potent immune escape mechanism ^57–59^. Given that HRD is a major driver of chromosomal instability (CIN), and harbor high levels of aneuploidy^60^, we hypothesized that CIN could contribute to immune escape in HRD tumors. One-third of their genomes exhibit LOH, and, as expected, the frequency of LOH is higher in HRD cases compared to HRPs (**Figure S5A, B**). Supporting this hypothesis, HRD ovarian cancers and triple-negative breast cancers (TNBCs) bear a higher rate of HLA LOH compared to HRP tumors of the same histologies^15,61^. This event is further exacerbated in patients with end-stage ovarian cancer^58^. However, the genome-wide profile of LOH, beyond HLA-loss, and its connection with immune escape remain largely unexplored.

To further understand how CIN can contribute to immune escape in HRD tumors, we compared LOH and amplification rates in our two most divergent tumor subtypes: dHpC and dCpH (**Figure 4A; Table S4**). Strikingly, HRD tumors within the dCpH group exhibited frequent LOH in genomic regions containing key immune regulators, particularly those involved in the cGAS/STING-dependent inflammatory signaling cascade. Loci on chromosomes 3p, 5q, and 9 had the highest frequency of LOH among regions enriched in dCpH tumor types (**Figure 4A, top; Figure S5A**). LOH on chromosome 9p, which encodes for IFN genes (including subunits of IFN-α and IFN-β), as well as *JAK2* and *CD274 (PD-L1)*, was detected in more than 75% of samples in dCpH HRD tumors compared to 55% in dCpH HRP. This locus has been previously implicated in immune escape in acute myeloid leukemia and in all the tumor types in our dCpG group (HNSC, LUAD/LUSC and SKCM-met)^59,62–66^. Additional LOH regions enriched in dCpH HRD tumors included chromosome 5q, which encodes *STING1*, along with other immune-related genes such as *IL12B* and *IRF1,* and chromosome 3p, which encodes *MYD88*, *CCR1*, and *CCR5*, all genes critical for immune sensing, inflammation, and leukocyte recruitment. In HRD tumors, the LOH of these loci frequently co-occurred with 9p LOH (*p* = 1.3e-10, OR = 3.24, and *p* = 3.0e-06, OR = 2.02, respectively for chr3p and chr5q loci, respectively), suggesting a concerted loss of key immune signaling genomic clusters in immune-evasive HRD cancers.

**Figure 4.**
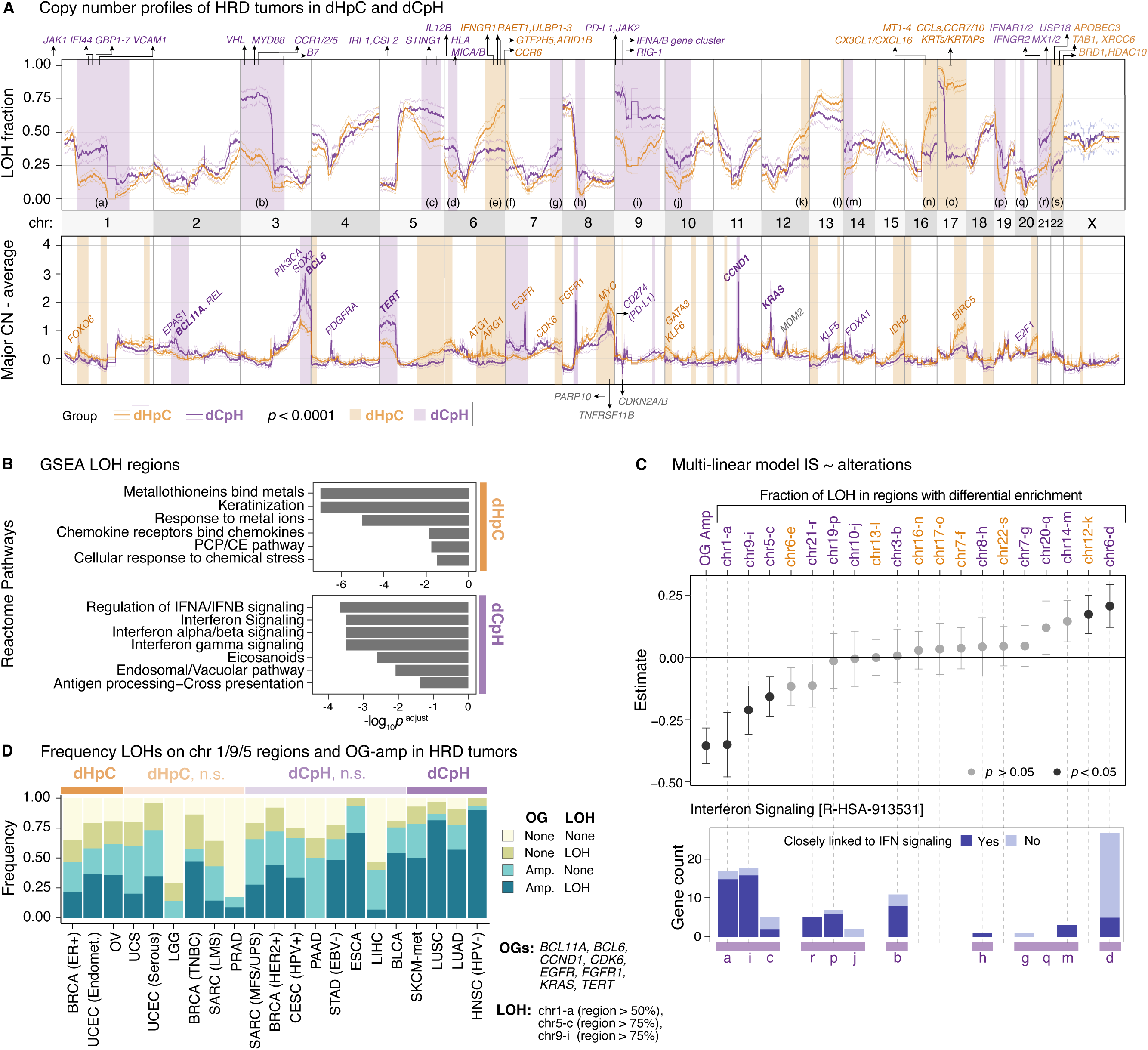
Immunomodulatory alterations in HRD tumors contribute to the tumor type-specificity of inflammatory responses. **(A)** Fraction of LOH (**top**) and average amplification profile (**bottom**) in HRD samples of the dHpC and dCpH categories are shown. Amplifications are determined as average values across samples after subtracting the copy-number baseline (mean chromosome-level major allele copy number in each sample). The genome was binned into 100 kb intervals; 95% confidence intervals for both variables were calculated by one-sample t-tests against the null hypothesis of zero mean difference. Selected genes in regions of interest are marked; large regions with significant differences are shaded, with colors indicating the group with higher alteration prevalence. X chromosome LOH is calculated using only the samples from female patients. **(B)** Reactome pathway enrichment analysis of genes in LOH regions differentially enriched in dHpC or dCpH; only pathways with adjusted p < 0.05 are shown. **(C) Top:** β-coefficients for covariates (LOH and oncogene amplification status) used in the multiple linear regression of Z-normalized IS. Oncogene amplification is defined as amplification in any of BCL11A, BCL6, CCND1, CDK6, EGFR, FGFR1, KRAS, or TERT. LOH status is defined as the fraction of LOH in regions marked in panel A (**Table S5**). **Bottom:** Number of genes per LOH region in the Interferon Signaling pathway enriched in dCpH LOH regions; antigen-presentation–related genes are highlighted in light blue. **(D)** Prevalence and co-occurrence of oncogene amplifications and LOH across tumor types, ordered as in Figure 2A.

To functionally characterize these recurrent genomic losses, we conducted pathway enrichment analysis using the Reactome database^36^ (**Methods**). Among the 4,760 genes (**Table S5**) located within LOH regions enriched in dCpH cancer types, we identified significant depletion of pathways involved in type I and II interferon signaling, and antigen processing and presentation (**Figure 4B; Table S6**), essential for mounting effective anti-tumor immune responses. In contrast, pathway enrichment analysis (**Table S6**) of the 2,554 genes (**Table S5**) residing in LOH regions enriched specifically in dHpC samples revealed an enrichment in the loss of pathways related to keratinization in dHpC tumors (**Figure 4B**). This may reflect a shift toward epithelial-to-mesenchymal transition (EMT), a cellular state associated with enhanced invasion and metastasis^67,68^. Additionally, we observed loss of pathways involved in the cellular response to metal ions (**Figure 4B**), which are known to accumulate in certain tumor types, including ovarian cancer^69,70^. Disruption of metal ion detoxification may contribute to a toxic tumor microenvironment detrimental to infiltrating immune cells. Interestingly, we also identified potential loss of “chemokine receptor binding to chemokines” driven by the recurrent loss of multiple *CCL* chemokine genes located on chr17q (**Figure 4B, C**). This event likely links with the HRD onsetting LOH of chr17q in tumors with *BRCA1* deficiency, which is more prevalent in dHpC tumor types such as breast and ovarian cancers (**Figure 4A**). Consistently, LOH of this region is more common in *BRCA1*^−/−^ compared to *BRCA2*^−/−^ HRD tumors (88.6% vs 36.0% had > 50% of the locus with LOH, respectively, Fisher’s exact test, p < 2.2e-16, OR = 13.7).

Together, these findings suggest that while elevated LOH is a universal feature of HRD tumors, the immunological consequences of LOH are tumor-type specific. In dCpH tumors, LOH preferentially disables interferon signaling and antigen presentation, promoting direct immune evasion. In dHpC tumors, LOH may instead drive EMT-related invasiveness, modulate immune cell recruitment, and create a metabolically hostile microenvironment *via* impaired metal ion detoxification, thereby contributing to alternative modes of immune remodeling.

In addition to the distinct LOH profiles, we observed pronounced differences in amplifications between the dHpC and dCpH tumor-type groups. Notably, amplifications of oncogenes (OG) such as *BCL11A*, *BCL6*, *CCND1*, *CDK6*, *EGFR*, *FGFR1*, *KRAS*, and *TERT* were significantly more prevalent in dCpH HRD tumors compared to dHpC counterparts (**Figure 4A**, **bottom**; **Figure S5B, C; Table S7**). These amplifications frequently co-occurred (**Figure S5D**) and were collectively associated with a reduced IS (**Figure S5E**). Accordingly, we defined OG amplified samples as the union of tumors harboring any of these alterations. Although OG-activating mutations were less frequent in HRD tumors compared to HRP (**Figure S5C**), we found that when present, mutations in *KRAS* and *NRAS* were enriched in immunologically cold-HRD tumors compared to hot-HRD, mirroring the amplification trends. This suggests that oncogenic stress, known to modulate anti-tumor immune responses^71^, may contribute to the observed tumor-type heterogeneity and to immune suppression or exclusion in dCpH HRD tumors.

Next, we extended our analysis to all differential LOH regions, 12 in dCpH and 7 in dHpC (**Figure 4A**, shaded areas), to determine which of these most strongly influences inflammation levels in HRD tumors. In pan-cancer HRD tumors, we built a multivariate linear regression model to predict IS (Z-score normalized within each tumor type) using LOH and OG amplification status as covariates (**Figure 4C**). This analysis revealed that LOH at the dCpH-enriched chr1 region (loci-a) was among the strongest negative predictors of IS, along with LOH at chr9-i and chr5-c loci, and the OG amplifications (**Figure 4C; top**). The chr1 LOH region harbors multiple immune-related genes such as *JAK1*, *IFI44,* and members of the GBP family genes on the p-arm, and spans the *CD1– IFI16* locus on the q-arm, which was previously identified as a frequent epigenetic silencing hotspot relevant to immune evasion in prostate cancers^72^. IFN signaling pathway genes (**Figure 4C**) were distributed across multiple differentially enriched LOH loci in dCpH, particularly in those predicted to have a high impact in reducing IS. To understand the broader applicability of these findings in pan-cancer settings, we examined the distribution of CIN-driven immune escape mechanisms across the full pan-cancer dataset (**Figure 4D**; tumor types ordered identically to **Figure 2A**). We observed a gradient of increasing immune escape frequency as we moved from dHpC to dCpH through intermediate tumor types not assigned to either extreme group. Tumors such as BLCA, ESCA, and STAD, classified as dCpH-n.s., displayed amplification and LOH profiles highly similar to dCpH, reinforcing the idea that these mechanisms extend beyond the initially defined extremes. The prevalence and combinatorial impact of LOH and oncogene amplifications may thus account for intermediate or attenuated inflammatory phenotypes in these tumors, helping explain heterogeneity in HRD IS profiles across histologies.

Collectively, these findings highlight that CIN-driven immune escape mechanisms via immune gene LOH and oncogene amplification are widespread in HRD tumors and play a determining role in determining their immune phenotype and may override the expected immunogenic consequences of HR deficiency.

### Chronic inflammation, senescence, and angiogenesis drive immune escape in HRD tumors

Given that immune evasion in HRD tumors may arise from both genomic alterations and transcriptional reprogramming, we performed integrative pathway enrichment analyses in HRD versus HRP tumors to systematically quantify these mechanisms and assess their immune modulatory consequences. We identified differentially expressed and genomically altered pathways enriched in HRD tumors, highlighting immune-modulatory programs shaped by both mutations and CNAs. Next, we correlated these enriched Reactome signatures with inferred immune cell proportions across tumor-type groups. This integrative approach (**Figure 5**) allowed us to delineate distinct immune escape strategies driven by genomic instability versus those shaped by inflammatory remodeling. In the following section, we focus on a curated subset of immune signaling and immunomodulatory pathways; the complete list of enriched pathways and their TME correlations is provided in **Table S8-10**.

**Figure 5.**
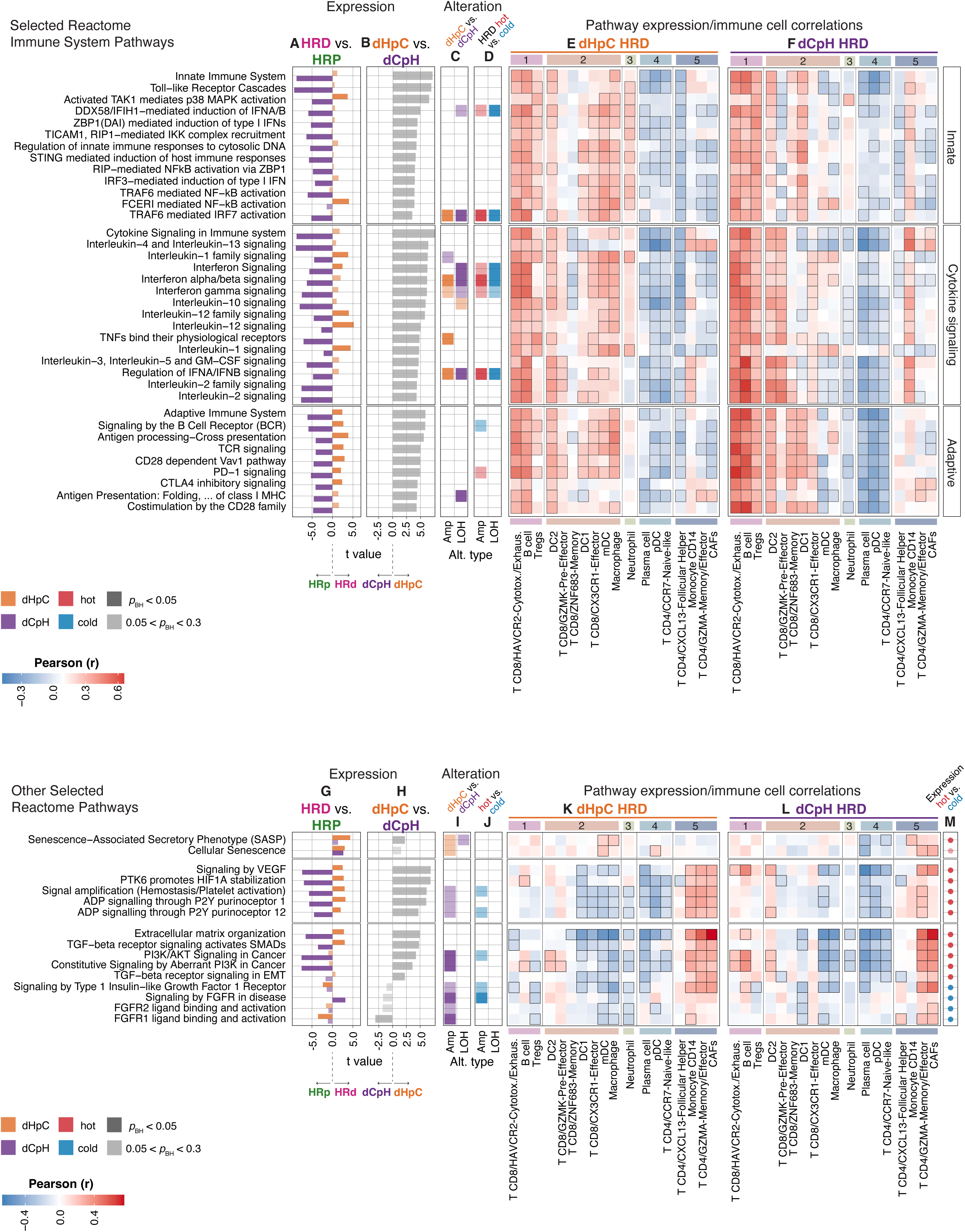
Angiogenesis and senescence as immune escape mechanisms prevalent in HRD tumors. **(A)** T values comparing Z-normalized Reactome pathway ssGSEA scores in HRD (positive t values) versus HRP (negative t values) tumors, analyzed separately in dHpC (orange) and dCpH (purple). **(B)** T values comparing Z-normalized ssGSEA scores between dHpC HRD and dCpH HRD. **(C)** Reactome pathway enrichment analysis of amplifications (2 copies above baseline major copy number) and LOH in dHpC versus dCpH HRD tumors; colors denote enrichment in dHpC (orange) or dCpH (purple). **(D)** Same as in panel C, comparing hot HRD versus cold HRD tumors. **(E)** Pearson correlation coefficients between Z-normalized ssGSEA scores and Z-normalized immune cell proportions (clusters as in Figure 3) in dHpC HRD tumors; significant correlations (p < 0.05) are outlined with black border. **(F)** Same as in panel E but for dCpH HRD tumors. **(G-L)** Additional pathways (in the same order as panels A–E). **(M)** T values from hot HRD versus cold HRD comparison; red indicates enrichment in hot and blue in cold tumors. For panels A-D, G-J, and M, the shades indicate the statistical significance (adjusted *p* < 0.05, darker shades).

To investigate immune signaling differences associated with HRD across tumor types, we identified differentially expressed pathways between HRD and HRP samples. The enrichment analysis was performed using Z-score normalized ssGSEA scores within each tumor type before combining them into dHpC and dCpH groups. This analysis revealed a broad downregulation in innate immune, cytokine, and adaptive immune signaling pathways (**Figure 5A**, **Figure S6**) with HRD in dCpH group. Compared to their dCpH counterparts, dHpC HRD tumors showed marked upregulation of interferon signaling, broad myeloid cell activation via RNA and DNA sensing cascades, and pattern recognition receptor signaling (**Figure 5A**, **Figure S6)**. This phenomenon co-occurred with elevated antigen presentation and cross-presentation, B cell and T cell receptor signaling, as well as inhibitory checkpoint pathways, including PD-1 and CTLA-4 signaling (**Figure 5A**, **Figure S6**). We also directly compared the normalized pathway expression in HRD tumors for dHpC and dCpH (grey bars), and dHpC tumors exhibited elevated expression of all major innate and adaptive immune pathways (**Figure 5B**, **Figure S6**).

Next, to link these transcriptional changes with underlying genomic drivers, we integrated copy number alterations with immune pathway activity. This revealed that type-I interferon, *TRAF6*-mediated *IRF7* activation, and TNF signaling pathways were recurrently amplified in dHpC and deleted in dCpH (**Figure 5C**). The enriched amplifications in dHpC suggest that tumor-intrinsic mechanisms are actively driving proinflammatory pathways, potentially paving the way towards chronic inflammation, a finding we previously documented in ovarian cancer^9^. Notably, this dichotomous amplification/deletion pattern also emerged when comparing hot-versus cold-HRD tumors across the full pan-cancer cohort, independent of tumor type (**Figure 5D**), therefore reinforcing the broader relevance of this mechanism. As a potential downstream consequence of these genomic alterations, dHpC HRD tumors exhibited clear transcriptional evidence of chronic inflammation, including elevated NF-κB activation (mediated in part by Myd88 and FCERI signaling, potentially resulting from myeloid cells as well as the upregulation of TGF-β-activated kinase 1 (TAK1)-mediated p38 MAPK activation (**Figure 5A, Figure S6**). Interestingly, among cytokine signaling pathways, interleukin-1 (*IL-1*) signaling stood out as highly upregulated, further supporting the proinflammatory and chronically inflamed phenotype of dHpC HRD tumors.

To gain a comprehensive view of how transcriptional and genomic alterations shape the TME, we correlated pathway-level expression with inferred immune cell proportions and integrated these associations with genomic and transcriptomic modalities. Across all tumor types, immune signaling pathways showed strong positive correlation with inferred proportions of cytotoxic/exhausted CD8⁺ T cells and B cells (in cluster 1), likely driven by antigen recognition and costimulation. In dHpC tumors, where these pathways are transcriptionally upregulated (**Figure 5A**, **Figure S6**), their correlation with cluster 1 cell types suggests coordinated enrichment of both inflammatory signaling and immune effector cells (**Figure 5A, E, Figure 3A**). In dCpH, where immune pathways were generally downregulated (**Figure 5A**, **Figure S6**), these same positive correlations (**Figure 5F)** imply a reduced abundance of effector cells in tandem with impaired signaling, as we previously demonstrated (**Figure 3A**). Beyond cluster 1, precursor and effector CD8⁺ T cell populations of cluster 2 were also positively correlated with immune signaling pathways, particularly in dHpC (**Figure 5A, E, F**). Myeloid populations (DC1, DC2, mDCs and macrophages) in cluster 2 displayed histology-specific patterns: in dHpC tumors, mDCs and macrophages positively tracked with inflammatory signaling **(Figure 5E**), whereas in dCpH, these associations were absent (**Figure 5F**), suggesting impaired myeloid activation. Interestingly, DC2 cells exhibited positive correlations with immune pathways in both dHpC and dCpH groups **(Figure 5E, F**), suggesting they may represent a more conserved or constitutive axis of myeloid involvement in HRD tumors, potentially involved in tolerogenic or transitional immune states.

The chronically inflamed phenotype of dHpC HRD tumors, marked by upregulated TAK1-mediated p38 MAPK activation and elevated IL-1 signaling, suggests enhanced cellular stress responses and engagement of the senescence-associated secretory phenotype (SASP). Notably, TAK1 (TGF-β–activated kinase 1) functions as an upstream activator of p38 MAPK via the MKK3/6 pathway and plays a pivotal role in SASP development^73^. To explore this further, we compared signal transduction pathway activity between HRD and HRP tumors. HRD tumors, particularly those in the dHpC group, showed significant upregulation of multiple senescence-associated pathways, with SASP emerging as the most prominently enriched (**Figure 5G,H; Figure S7A**). In dHpC HRD tumors, cellular senescence and SASP induction were driven by both transcriptional upregulation and further supported by a moderate enrichment of copy number amplifications (e.g., *CEBPB*, *JUN*, and *VENTX*) (**Figure 5I**). dHpC samples were characterized by overexpression of canonical inflammatory mediators such as *IL6*, *IL1A/B*, and *CCL2/3* **(Figure S7A, B**). In the hot samples among dHpC HRD cases, *STAT3, VENTX*, and *NFKB1* were further upregulated, suggesting a potential path toward chronic inflammation driven by SASP^74–76^ even in the inflamed subset. Moreover, the SASP program correlated positively with mDCs and macrophages in dHpC HRD tumors (**Figure 5K**). In hot-HRD tumors (**Figure 5M**), this inflammatory SASP may act as a double-edged sword: sustaining immune activation while simultaneously driving immune dysfunction and the polarization of immune cells into immunosuppressive states^77–86^.

In contrast, dCpH HRD tumors exhibited a distinct cellular senescence profile characterized by the upregulation of growth and cell-cycle-related factors, including core histone genes, the APC/C complex, and *IGFBP*s-features (**Figure S7A, B)** that are associated with oncogene- or growth factor-induced senescence (**Figure 5G, H).** Cellular senescence pathway expression positively correlated with CAFs and CD4⁺ effector/memory T cells (**Figure 5L**). In the dCpH group, oncogene amplifications that resulted in upregulation of pathways related to FGFR signaling have a similar impact on the TME of dCpH group (**Figure 5K, L).**

Consistent with a chronic inflammatory state, VEGF signaling and VEGFA expression were upregulated in HRD tumors (**Figure 5G**; **Figure S7C**), expanding prior findings in BRCA1-deficient murine ovarian models to a broader spectrum of HRD and gynecological cancers ^9,87^. Importantly, this angiogenic program appeared to diverge between tumor types: in dHpC tumors, VEGFA upregulation co-occurred with inflammatory gene expression, while in dCpH, it was coupled with oncogene activity (**Figure S7C**), suggesting distinct regulatory mechanisms. The dHpC-specific pattern aligns with our previous reports implicating cGAS/STING signaling in the regulation of VEGFA in HRD tumors^9^. Adenosine signaling pathways^88^ were also preferentially activated in dHpC HRD tumors (**Figure 5G**), marked by transcriptional upregulation of CD38 and ADORA1 (**Figure S7D**). Notably, these pathways lacked corresponding copy number amplification in dHpC (**Figure 5I, J**), suggesting potential transcriptional activation that may also be modulated by the TME.

We examined how these mechanisms correlate with TME cell populations. Angiogenesis, adenosine, and TGF-β signaling, extracellular matrix organization pathways, which are upregulated in dHpC, correlated positively with cell proportions of cluster 5, including CAFs, CD4^+^ memory/effector and follicular helper cells, and monocytes (**Figure 5K, L**), reinforcing a close link between angiogenesis and immune cell exclusion.

Together, these findings reveal that HRD tumors engage distinct immune escape strategies shaped by the interplay of chromosomal alterations, transcriptional rewiring, and tumor-type-specific microenvironmental cues. In dHpC cancers, HRD drives a chronic inflammatory state marked by interferon and cytokine pathway amplification, inflammatory SASP, angiogenesis, and adenosine signaling, all features that correlate with robust immune infiltration but also exhaustion and immune dysfunction. Conversely, dCpH tumors exhibit widespread LOH of immune-regulatory loci, reduced immune signaling, and oncogene-driven senescence, promoting a more stealth-like immune evasion. These divergent immune programs underscore the need for tailored immunotherapeutic strategies that consider both tumor-intrinsic and extrinsic determinants of immune visibility in HRD cancers.

## Discussion

HRD is a defining feature of a clinically actionable subset of tumors that respond to DNA-damaging agents and PARP inhibitors. Beyond its role in synthetic lethality, HRD has been proposed as a determinant of tumor immunogenicity, yet the pan-cancer immune consequences of HRD remain poorly understood.

Our study delineates a pan-cancer immune taxonomy of HRD tumors, uncovering mechanistically distinct immune programs with implications for PARPi combinations. While prior research has largely focused on gynecologic cancers^15,16,89^, we broaden the scope by applying a novel whole-exome sequencing classifier *CSI-HRD* to >10,000 TCGA tumors and systematically characterizing immune profiles across 33 cancer types.

The findings reveal that HRD is not uniformly associated with immune activation but rather underlies two distinct immune states shaped by tumor context and chromosomal instability. We uncover two immune archetypes among HRD tumors: those with elevated inflammation (deficient-Hot/proficient-Cold; dHpC) and those with suppressed immune activity (deficient-Cold/proficient-Hot; dCpH). dHpC tumors, including ER^+^ breast, ovarian, and endometrial cancers, show enriched infiltration of cytotoxic CD8⁺ T cells, Tregs, and myeloid subsets, alongside robust IFN and cytokine signaling. By contrast, dCpH tumors, such as lung, melanoma, and head and neck cancers, exhibit dampened immune signaling, depletion of tumor-reactive lymphocytes, and marked evidence of immune evasion.

Multi-omic integration reveals that these phenotypes are shaped by tumor-intrinsic and extrinsic mechanisms. dHpC tumors amplify innate immune pathways and display transcriptional features of chronic inflammation, including NF-κB and SASP activation, adenosine signaling, and VEGF-driven angiogenesis. These programs co-occur with CD8⁺ T cell exhaustion and M1 macrophage polarization, suggesting a double-edged role in promoting and restraining immunity. Conversely, dCpH tumors harbor widespread LOH at immune regulatory loci, including JAK-STAT, *STING1*, and *IFNA/B* gene clusters, alongside frequent oncogene amplifications. These genomic alterations correlate with suppressed inflammation, reduced myeloid activation, and immunologically inert TMEs. Surprisingly, HLA LOH emerged as a positive predictor of inflammation in multivariate models, potentially reflecting immune editing in tumors with pre-existing cytotoxic infiltration.

These findings offer a new lens to interpret immunotherapy responsiveness in HRD cancers. dCpH tumor types already benefit from approved ICB options. However, dHpC tumors, despite being enriched for inflammatory signatures, respond poorly to ICB in clinical trials. Our results suggest that stratifying patients by HRD and immune context^10^ may help identify candidates most likely to benefit from ICB plus PARPi or triple combinations including anti-angiogenic agents^9^. Importantly, we find that a subset of dCpH tumors exhibits hot-HRD features with elevated SASP, angiogenesis, and adenosine signaling. These may represent escape-prone subpopulations for which rational combinations targeting chronic inflammation^90–92^ and immune suppression *via* the adenosine pathway, SASP ^93,94^, and exhaustion could prove effective.

While our pan-cancer RNA-based immune cell deconvolution provides broad insights and our sample size allows generalizable insights across histologies, it lacks the resolution of single-cell and spatial profiling. Moreover, we do not assess clinical response to PARP inhibitors or ICB directly, and our hypotheses regarding therapeutic benefit require validation in prospective trials stratified by HRD and immune context. Future work should prioritize single-cell, spatial, and clinical trial-linked analyses to validate these immune programs as predictive biomarkers.

In conclusion, our multi-layered analysis enabled us to identify how specific immune signaling and evasion mechanisms co-occur with distinct patterns of immune cell infiltration or exclusion, offering deeper insights into the interplay between tumor-intrinsic alterations and microenvironmental remodeling. It revealed that HRD is not a uniform predictor of tumor immunogenicity. Instead, chromosomal instability, immune gene dosage, inflammation, and stromal remodeling interact to dictate immune visibility. This immune bifurcation of HRD tumors provides a blueprint for refining biomarker strategies and combinatorial immunotherapies targeting HRD tumors.

## STAR Methods

### Key Resources Table

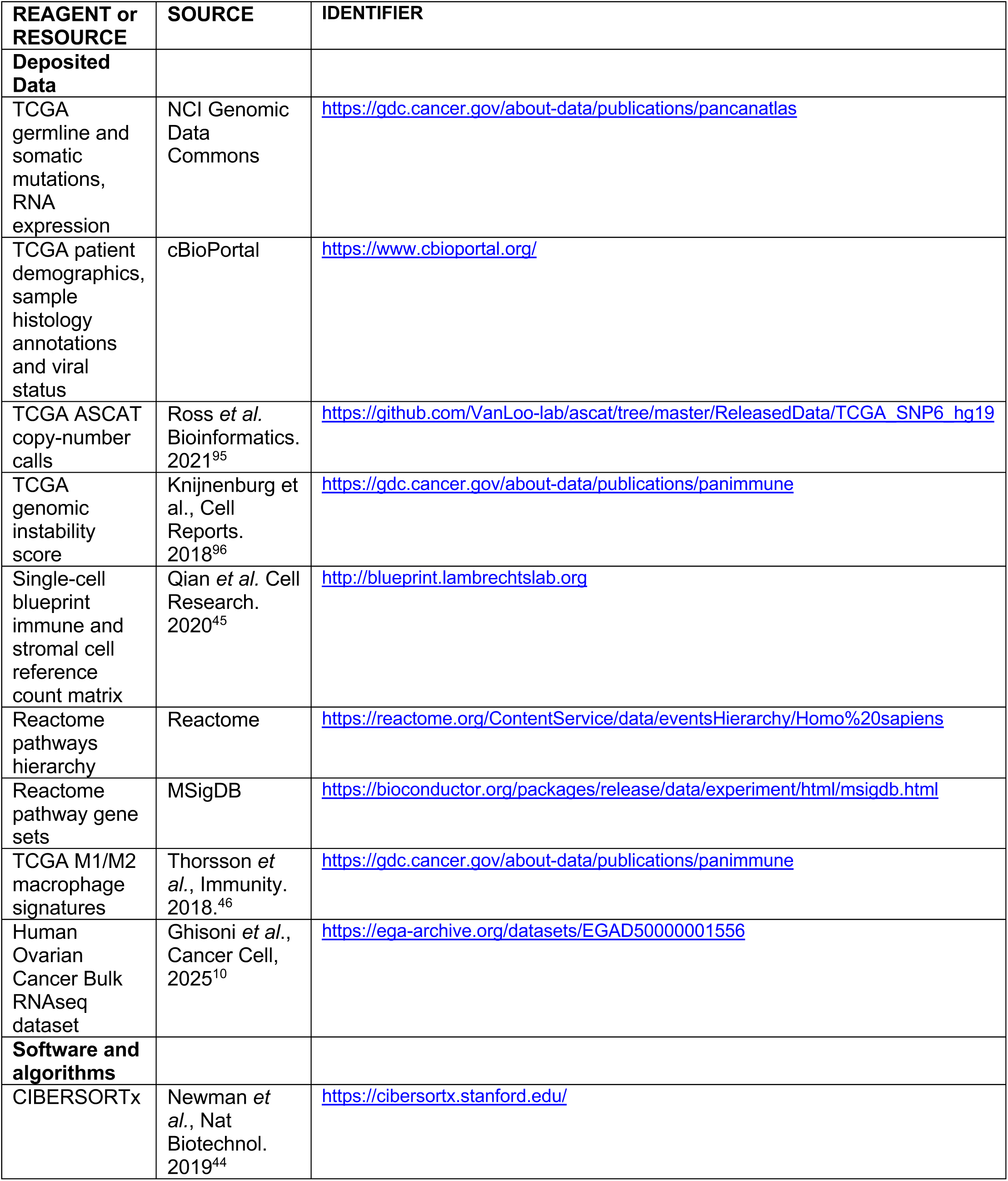
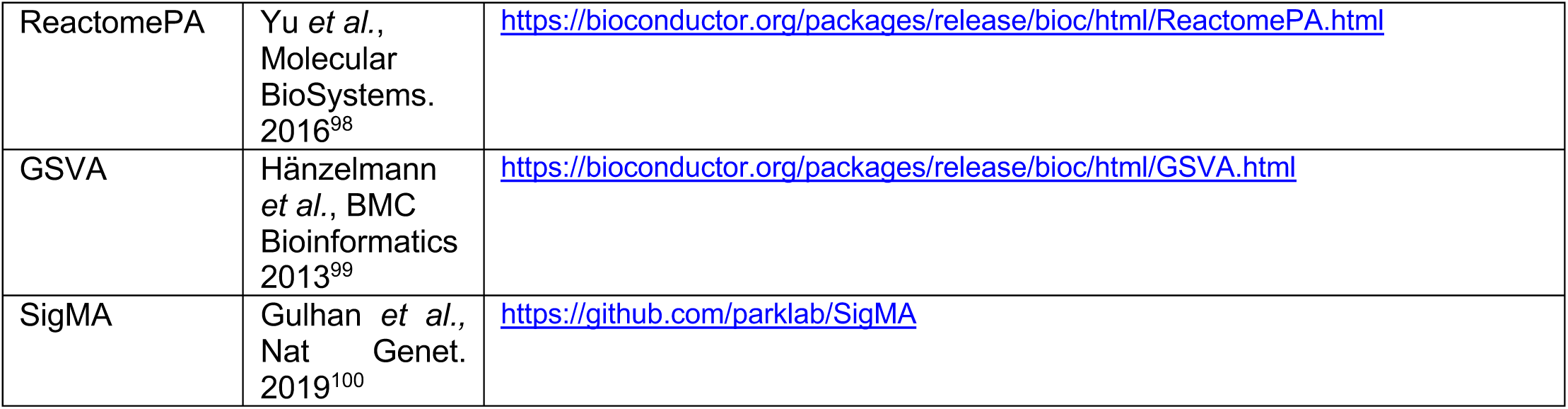

### Method Details

#### Samples excluded from analysis

The 91.8% of TCGA data has both WES and gene expression data. The 97.8 only has WES and 92.1 only gene expression. As a result, for these samples we cannot calculate the inflammation score and HRD status simultaneously, and they are excluded from the parts of analysis that includes information of both the inflammation and genomic properties. Tumors with high 8-oxo-guanine related artefactual mutations (94/10308, 0.9% of all samples) as reflected by presence of SBS45 were removed from the analysis.

#### Gene expression

We used the pre-processed RNA sequencing gene expression file ([EBPlusPlusAdjustPANCAN_IlluminaHiSeq_RNASeqV2.geneExp.tsv](http://api.gdc.cancer.gov/data/3586c0da-64d0-4b74-a449-5ff4d9136611)) from Pan-cancer atlas repository. For deconvolution analyses using CIBERSORTx^44^, we used the TCGA counts matrix that was assembled manually by concatenating individual cancer type counts data taken from the GDAC firehose database.

#### Somatic mutations

The processed somatic mutation data was taken from the Pan-cancer atlas repository ([mc3.v0.2.8.PUBLIC.maf.gz](http://api.gdc.cancer.gov/data/1c8cfe5f-e52d-41ba-94da-f15ea1337efc)) which calculated by the MC3 collaboration aiming at a consensual and unified variant calling^34^. We considered as somatically mutated if one of the following holds:

- The mutations were truncating (“Nonsense_Mutation”, “Frame_Shift_Del”, “Frame_Shift_Ins”, “Translation_Start_Site”, “Splice_Site”)
- If the nonsynonymous mutations (“Missense_Mutation”, “In_Frame_Del”, “In_Frame_Ins”,“Nonstop_Mutation”) were “deleterious” in the SIFT field and “probably_damaging” in the PolyPhen.
- If the nonsynonymous mutations were predicted to be “pathogenic” or “likely_pathogenic” by ClinVar^97^.
- In any case of the cases above if a mutation was labelled as “Benign” according to ClinVar we did not consider this mutation.
- *Tumor mutational burden:* Tumor mutational burden was calculated as the rate of nonsynonymous mutations within 1 Mbp SNVs and indels.

#### Germline mutations

Germline mutations were processed in Huang et al.^101^ and obtained from GDC data portal (PCA_pathVar_integrated_filtered_adjusted.tsv), and the variants of unknown significance were excluded.

#### Copy-number alterations

Allele-specific copy number calls for TCGA samples were obtained from the ASCAT release on GitHub (https://github.com/VanLoo-lab/ascat/tree/master/ReleasedData/TCGA_SNP6_hg19). We evaluated copy number changes at four levels: (i) sample, (ii) chromosome, (iii) gene, and (iv) genome-wide in 100 kb bins. For each chromosome, we computed segment length–weighted averages of minor and major copy number values using all segments larger than 100 kb. A sample-level weighted average was then obtained by aggregating across chromosomes. Total copy number was defined as the sum of the minor and major averages.

##### Gene-level analysis

Copy number gains of 2 or 3 copies above the sample-level baseline were classified as relative amplifications. Focal amplifications were defined as segments with log10(length) < 6.5 and at least two additional copies over baseline. Loss of heterozygosity (LOH) was defined as regions with a minor copy number < 1.

##### Genome-wide profiles

The genome was divided into 100 kb bins, and segment length– weighted averages of overlapping segments were used to compute minor and major copy number. LOH segments were designated where the minor copy number < 1. Amplification profiles were derived from deviations in major copy number relative to the sample-level baseline, thereby reducing the confounding effect of prevalent LOHs. Total copy number was not used, as it is strongly influenced by LOHs in homologous recombination–deficient (HRD) samples, and our aim was to keep LOH and amplification distributions as independent as possible.

##### Statistical analysis

For groups of samples, the fraction of LOH in a region was calculated, and significance of enrichment was assessed using t-tests against the null hypothesis. Amplification profiles were determined as the average of baseline-corrected major copy number values across samples; t-tests were used to assess shifts relative to the null (equivalently, shifts relative to the sample-level baseline). When comparing two groups, t-tests were performed on both the LOH fraction distributions and the baseline-corrected amplification distributions.

##### Sex chromosomes

The Y chromosome was excluded from all analyses. LOH for the X chromosome was assessed only in female patients.

#### Methylation data

RESET^102^ was run on the latest data freeze of the TCGA sample cohort^103^ to identify functional silencing DNA methylation events. RESET v1.1.1 was downloaded from the BitBucket repository (https://marcomina85@bitbucket.org/cso_repo/reset.git). The RESET process is split as follows: Data preprocessing is required to select, out of the 450k probes, those mapping to gene promoters and showing a stable behavior in normal tissue. To this end, only probes mapping to gene promoters (henceforth referred to as *promoter probes*) according to the FANTOM5 annotation were retained, as done in the original RESET publication. The promoter probes were further filtered by removing those showing high variance or intermediate methylation scores (mean > 0.1 and < 0.8) across normal tissue samples. The preprocessing was repeated separately for each tumor type, and results merged together by taking a consensus set of probes (only the promoter probes passing the filtering step in every tumor type were retained). The last step of RESET analysis checks whether the expression of gene *x* decreases in the cancer samples with hyper-methylation events in the promoter regions of gene *x*. If this is the case, the hyper-methylation event is called a functional silencing event. Finally, a sample-level filter is applied to require each silenced gene through hyper-methylation to have a reduced expression by 1 standard deviation of its value in the corresponding tumor type.

#### Patient demographics, cancer type and subtype annotations

The age, sex and cancer type were obtained from Pan-cancer Atlas Repository (TCGA-CDR-SupplementalTableS1.xlsx). We analyzed data from 32 cancer types, further subdividing them into clinically relevant categories. Stratifications were based on viral status (CESC, HNSC, LIHC, STAD), molecular or histological subtypes (BRCA, SARC, UCEC), or metastatic properties (SKCM). HPV status in HNSC and EBV status in STAD were obtained from the cBioPortal PanCancer Atlas clinical data. For LIHC, HBV and HCV annotations, as well as CESC HPV annotations, we used the supplementary materials from the corresponding TCGA molecular characterization studies^104,105^. SKCM was classified as primary versus metastatic, and UCEC as endometrioid versus serous, based on PanCancer Atlas annotations (downloaded from cBioPortal). BRCA tumors were divided into HER2+, ER+, and triple-negative (TNBC) groups using cBioPortal clinical annotations (Firehose and TCGA publication^106^). Specifically, HER2+ samples were identified first, and the HER2– group was then separated into ER+ and TNBC subtypes.

#### Inflammation score and inflammation status

We assigned an inflammation score using the bulk RNA-seq data to every tumor as follows: We first computed single-sample GSEA (ssGSEA scores) for four different signature scores reflecting various aspects of immune activation: (1) a double-strand break interferon signature from Bruand *et al*.^9^ (2) a chemokine gene signature that results in intratumoral T-cell recruitment from Dangaj *et al*.^8^ (3) a single-cell transcriptomic-derived signature of activated CD8 T cells from breast cancer tumors published by Azizi *et al*.^38^ and (4) a curated interferon-gamma signature taken from the Hallmark collection of the MSigDB database^98^. We calculated the per-sample average of four ssGSEA scores to obtain the Inflammation Score (IS). The inflammation status of each sample was defined using tertiles of IS (hot: IS > T2, intermediate: T1 < IS < T2, cold: IS < T1). This calculation was performed separately in each tumor type, using tumors without viral infection in LIHC (HCV/HBV), HNSC (HPV), CESC (HPV) and STAD (EBV) tumor types and without MMRD or polymerase proofreading deficiency-related hypermutations. For tumors that fall into one of the excluded categories in the tertile calculation, we still used the thresholds obtained in the remaining samples in that tumor type. At the tertile calculation breast cancer cohort was not yet separated into TNBC and non-TNBC subsets (non-TNBC group including HR^+^ or HER2^+^ breast cancers, were shortly referred to as BRCA in the manuscript), similarly a single tertile calculation was performed in melanomas without stratifying metastatic and primary cases. When differential IS analysis is performed in a combined set of samples including more than one tumor type, we first applied a Z-score transformation.

To validate our IS, we applied it to an independent cohort of immune-classified treatment-naïve ovarian cancers for which we had previously generated both bulk RNA-seq and high-resolution whole-slide immunofluorescence (IF) imaging of CD8^+^ tumor-infiltrating lymphocytes (TIL)^10^.

#### Genomic Instability Score

The genomic instability score (GIS) was computed as the sum of telomeric allelic imbalance (TAI), large-scale transition (LST), and loss-of-heterozygosity (LOH) by Knijnenburg *et al.*^96^, and deposited on GDC by Thorsson *et al.*^46^.

#### Homologous Recombination Deficiency status definition (CNA, SBS, and Indel-based HRD Classifier/CSI-HRD)

HRD classification was performed following a four-step procedure (**Figure 2A**).

##### 1. MMRD and POLE-exo sample classification

We developed multi-class classifier built-in in SigMA (v2.0)^107^ to classify TCGA samples into MMRP, MMRD or POLE-exo mutant classes. This classifier combines SBS signatures and MSISensor scores (obtained from Thorsson et al.^46^) improving upon the strategies using only MSI scores^107^. MMRD and POLE-exo samples were not considered for HRD classification in the next step. Our findings suggest that HRd in MMRd and POLE-exo tumors is rare, we observed 22 cases out of 444 with BRCA1-/- and out of those all apart from two with GIS = 72 all had low GIS (< 11). However, because it would not be possible to distinguish the effect of HRD on immunogenicity in these samples that are hypermutated, we do not consider them as HRD tumors in downstream analysis. For the remaining MMRP samples, we calculated single base substitution (SBS) signature 3 (Sig3) score with SigMA^31^ optimized for TCGA-MC3 mutation calls as described below.

##### 2. Sig3 predictions from SigMA

The single base substitution signature 3 (Sig3) was calculated using the SigMA algorithm^31^, which uses whole-genome sequenced (WGS) data as a reference to improve Sig3 detection from WES and gene-panels. Because TCGA-MC3 calls differed from the mutation calls from Broad GDAC Firehose that we used in our first publication^31^, we optimized new classifiers specifically for TCGA-MC3, for which we regenerated WES simulations to match the average mutation counts per tumor type. The WGS data is used to determine the average mutational spectra in tumors with Sig3, which we refer to as Sig3+, across different tumor types. These spectra provide the expected probability distribution for calculating the likelihood of Sig3 for a new sample. WGS data is also used to simulate smaller sequencing platforms from which several signature-related features (likelihoods, cosine similarities, signature exposures by non-negative least squares decomposition) are calculated and finally, using the Sig3 status in WGS data as truth tags and signature features we train multivariate (MVA) machine learning classifiers. In this study, we used SigMA for single base substitution mutations from WES TCGA data, and the SigMA algorithm was updated by analyzing 3476 WGS samples for this manuscript and includes machine learning classifiers for 18 more tumor types compared to our previous publication^98^, allowing comprehensive pan-cancer detection of HRD. These classifiers are publicly available and can be accessed using *data = ‘tcga_mc3’* setting in the *run* function. For most tumor types (28/33) WGS data was available, allowing the use of multivariate analysis to predict Sig3. For 23/28 tumor types, we supplemented the Sig3+ samples in that tumor type with Sig3+ samples from other tumor types after accounting for tumor type-specific differences in mutation counts, as discussed in our previous publication. For few tumor types (5/28, LAML, TGCT, THCA, THYM, UVM, PCPG) which lack a sufficient amount of corresponding WGS data, we used the *tumor_type = ‘other’* setting in running SigMA. For these tumor types, the MVA classifiers were not trained due to a lack of WGS data, so instead, the Sig3 exposure and Sig3 likelihood were used to calculate Sig3+ samples.

##### 3. Indels linked with HRD

We calculated the total count of deletions of size greater than or equal to 5, further stratifying them based on whether they overlapped with a microhomology of size larger than or equal to 2 or not, receiving contributions predominantly from DSB repair by microhomology-mediated end joining and non-homologous end-joining, respectively.

##### 4. HRD classifier

Next, we trained a pan-cancer gradient boosting HRD classifier by training a gradient boosting classifier (GBC) that uses Sig3 score from SigMA, GIS and deletions at microhomologies as features. We used this classifier to predict HRD status of all samples in the TCGA data. We trained a gradient boosting classifier (*gbm* R package) using as features: (i) SigMA Sig3 scores, likelihoods, and Sig3 exposure, (ii) GIS, and (iii) indel counts.

We trained the classifiers following two strategies (**Figure S1A**): (i) for each tumor specifically and (ii) a single one for the entire pan-cancer data. For each of these approaches, we performed training in two steps by updating truth labels based on the results of the first step. In the first step, we labelled samples with bi-allelic *BRCA1*/*2*^−/−^ as HRD to be used in training. These samples are supplemented with *BRCA1*/*2*^WT^ cases, selecting the top 3 highest similarity in the feature space, which increased the positive training set size by 34%. The size is not tripled, as we allowed each HRP sample to be selected more than once, but kept only one instance in the training. We labelled the rest of the samples as true HRP cases for the first training. We optimized parameters of the classifier using 5-fold cross-validation to minimize overfitting. In the next step, we relabeled previously HRP samples with a high score as HRD and vice versa, which more than doubled the HRD-labelled samples. Then a second classifier was trained in the same manner as the first step with 5-CV for parameter optimization. Note that similar iterative approaches using BRCA1/2-/- samples as a true training set have been followed in developing HRD classification methods from WGS^99^.

For tumor types with > 10 HRD training samples (BLCA, BRCA, CESC, ESCA, HNSC, LGG, LIHC, LUAD, LUSC, OV, PAAD, PRAD, SARC, SKCM, STAD, UCEC) we trained a tumor-type specific classifier. We pooled all other tumor types together and treated them as a single group training a classifier just for them. We also trained pan-cancer classifier using all tumor types without selection. Our final classification is determined based on tumor type-specific and pan-cancer scores as shown in **Figure S1**.

### Cell population deconvolution from bulk RNA sequencing

We used *CIBERSORTx*^44^ in order to deconvolute TME cell-type populations from the TCGA bulk RNA sequencing data. As a reference for the cell populations to deconvolute, we built a TME signature matrix using recently reported pan-cancer single-cell transcriptomics data^45^. We downloaded the expression count data from the Lambrechts lab website (http://blueprint.lambrechtslab.org) and merge them into a unique Seurat object (using the *Seurat* R package). We labeled the cells using the fine cell type calls kindly given by the authors upon request then subset the Seurat object in order to have a maximum of 400 cells per cell type. The matrix of count data was used as an input in *CIBERSORTx* to build a gene expression signature matrix (by disabling the quantile normalization parameter, other parameters set to default). We then run the deconvolution in absolute mode using the TCGA RNA sequencing count data as mixture matrix (500 permutations, disabled quantile normalization). The relative mode could be rescued by summing all populations to 1. We then compared the cell population frequencies or abundance by performing comparisons between different groups.

### Quantification and Statistical Analysis

The statistical tests performed, significance of the results are reported in the main text and methods discussed above as well as on the figures and their legends.

### Data and Code Availability

The TCGA data (dbGaP accession phs000178) is obtained through NCI Genomic Data Commons, cbioportal and previous publications as detailed in the Key Resources Table. Bulk RNA-seq data and mIF based immune categorizations for ovarian cancers used to validate inflammation score was obtained from our recent publication^10^ and deposited under at EGA (EGAD50000001556) and GEO (GSE264660). Any additional information required to reanalyze the data reported in this paper is available from lead contacts upon request.

## Acknowledgements

This work was supported by grants to DDL (Ludwig Institute for Cancer Research, DoD OC210038), to DCG (DoD OC230088), and PJP (NIH R01CA269805).

## Supplementary Figures

**Figure S1.**
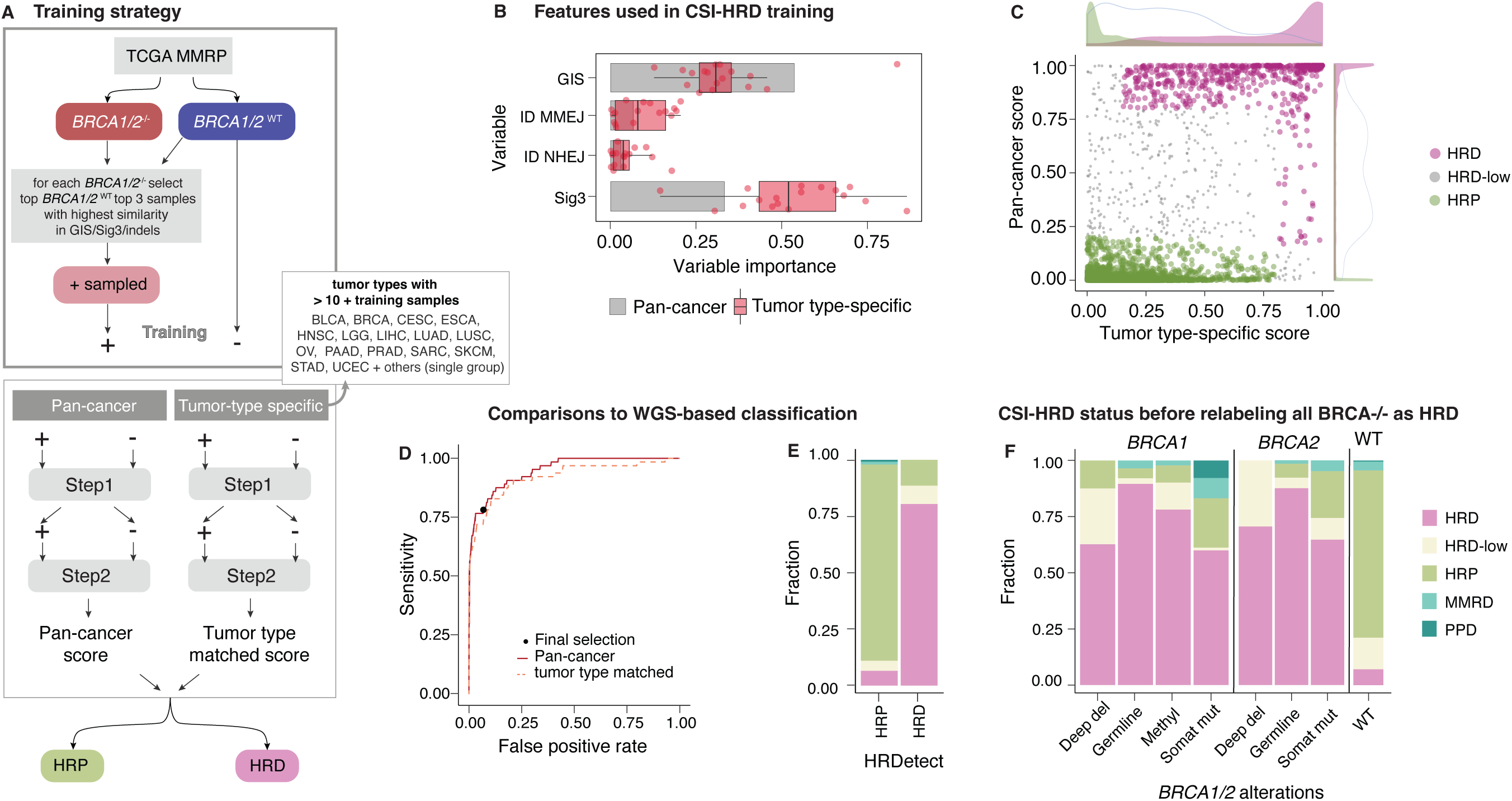
CSI-HRD classification strategy and performance. **(A)** Steps of the training procedure for the CSI-HRD training. As the first step, *BRCA1*/*2*^−/−^ samples were identified and used to select additional *BRCA1*/*2*^WT^ samples with matching Sig3, GIS, and indel profiles (three top matches picked for each *BRCA1*/*2*^−/−^ sample, with replacement, increasing the positive set overall by a factor of 2.3), while the rest of the samples were used as negatives. Two approaches were followed for training: (i) pan-cancer and (ii) tumor type-specific (for types with more than 10 positive samples, as listed). For the remaining tissues, all samples were combined. Both approaches followed a two-step procedure, with an initial classification and retraining using updated labels—mainly to account for additional HRD samples that may have been left in the negative group. **(B)** Variable importance of the four features that were combined into a multivariate score by the gradient boosting classifier. The bar plot shows values for the pan-cancer model; boxplots and data points indicate individual tumor type-specific classifiers. **(C)** Scatter plot showing CSI-HRD scores for pan-cancer and tumor type-specific classifiers. Colors indicate final categories (MMRD and PPD samples have been excluded). The density distributions on the top and right show projections of the scores on the two axes: filled colors represent HRD and HRP groups, and the HRD-low group is shown with a grey line. **(D)** Receiver operating characteristic (ROC) curves for pan-cancer and tumor type-specific scores (merging all tumor types). The black data point reflects the sensitivity and false positive rate for the selection outlined in panel C. **(E)** Fraction of samples in HRP, HRD, HRD-low, MMRD, and PPD categories—based on CSI-HRD from WES data—is shown within HRD and HRP groups defined by HRDetect WGS classification. **(F)** Fraction of HRP, HRD, HRD-low, MMRD, and PPD samples in BRCA1/2^−/−^ (subdivided by germline, somatic, deep deletion, or promoter hypermethylation events) versus BRCA1/2^WT^ samples.

**Figure S2.**
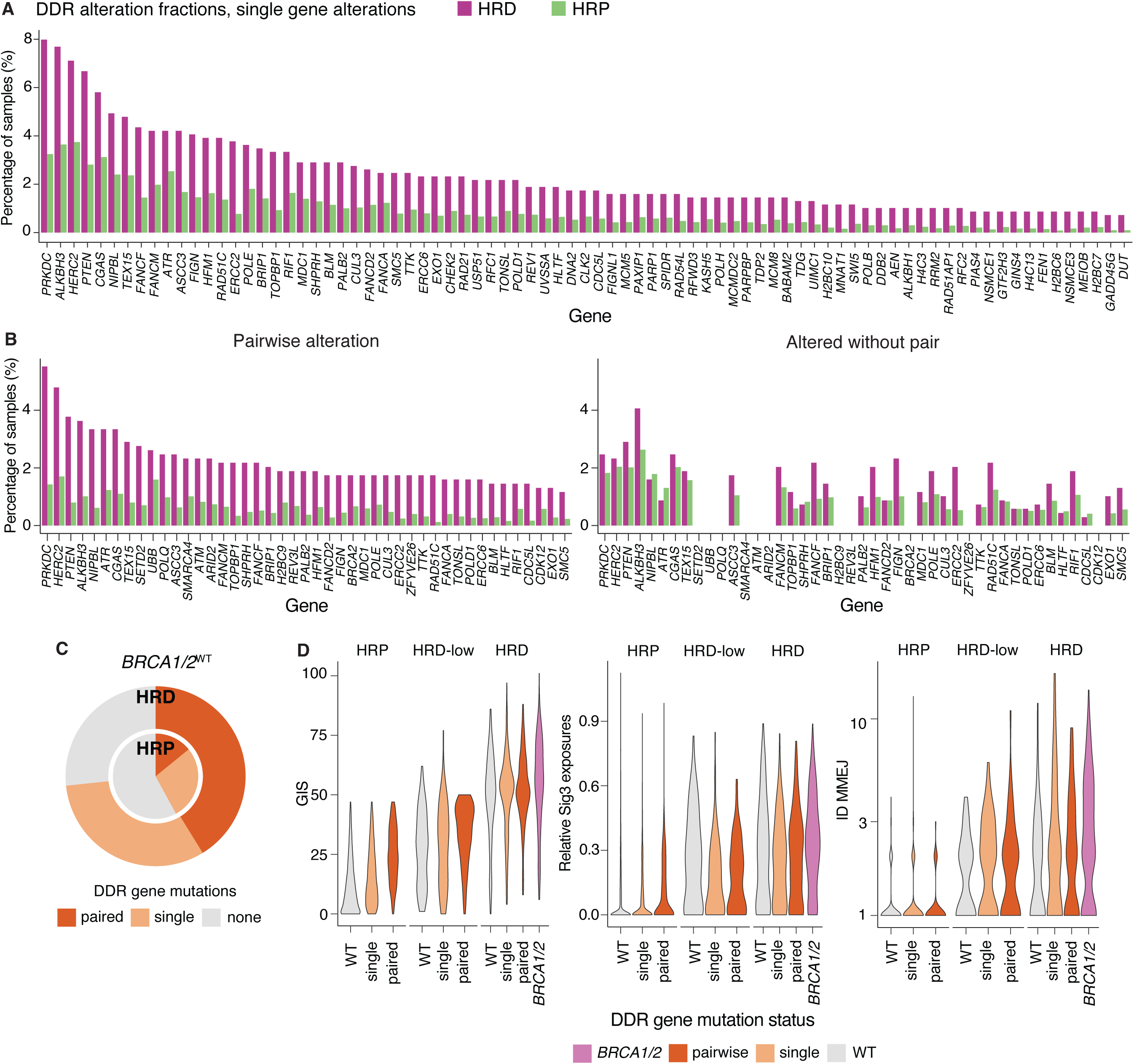
DNA damage repair (DDR) gene alterations enriched in HRD samples. **(A)** Samples with damaging mutations, deep deletions, promoter hypermethylation, or biallelic loss (due mutations + LOH, two mutations, deep deletion, or promoter hypermethylation) in DDR genes (Table S2) were identified. The prevalence of these alterations was compared (comparisons done for each alteration type) between HRD and HRP samples, and the bar plot shows the percentage of samples for genes with significant enrichment (all mutation types with significant enrichment combined into a broad alteration status). **B)** Instead of requiring only one DDR alteration, samples with a TP53-co-occurring pair of alterations (i.e., both present) were considered. The fraction of samples with a mutation in the presence of a significantly enriched pair (left), and those without such a pair (right), is shown. **(C)** Pie chart comparing the fraction of HRD and HRP samples with DDR alterations, further stratified based on whether they appear as pairs or as single alterations. **(D)** From left to right, the distributions of GIS, relative Sig3 exposure (exposure divided by total SNV counts), and ID counts at microhomologies are shown for samples stratified based on DDR alteration status and HRD status.

**Figure S3.**
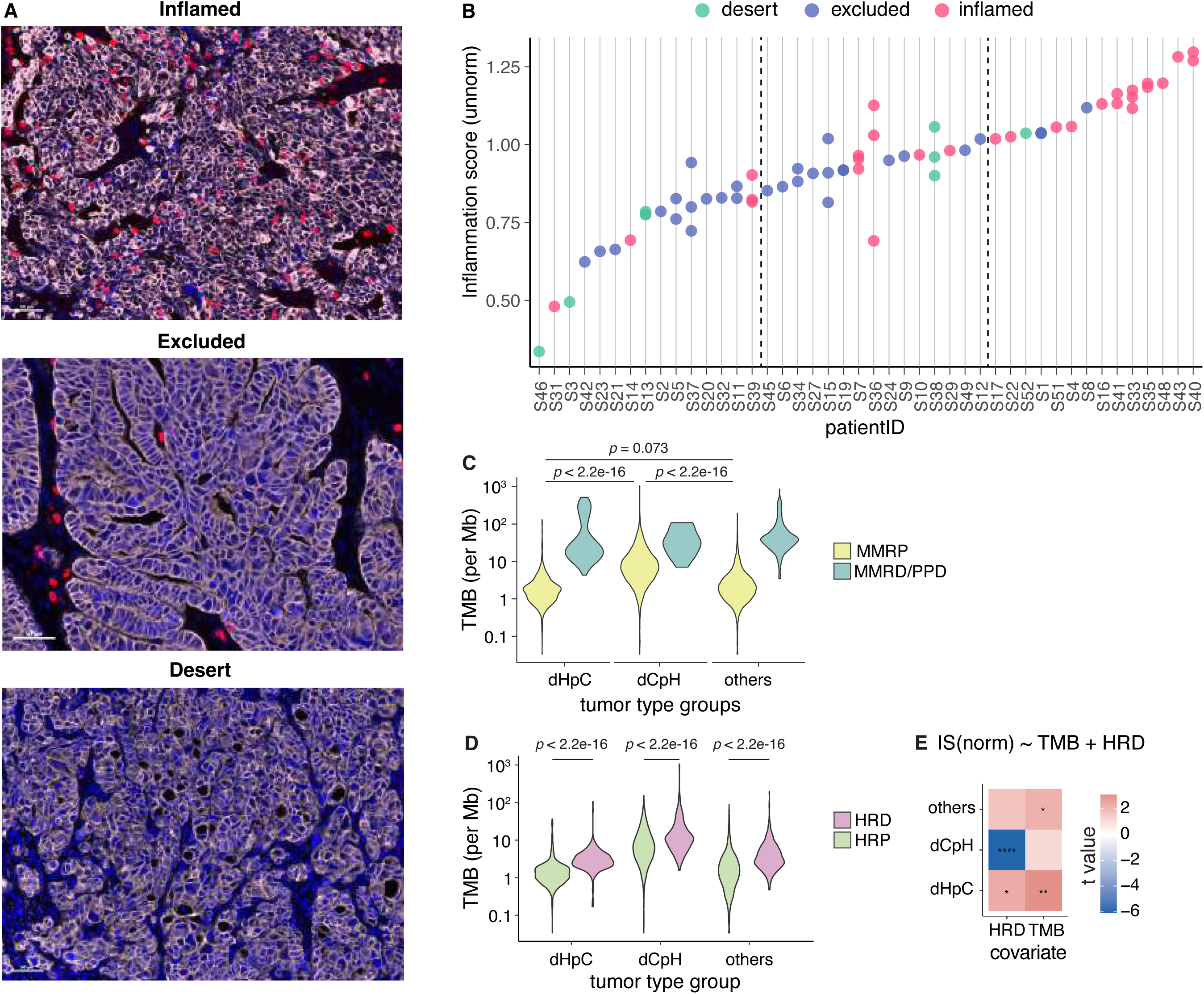
Validating the inflammation score and assessing the confounding effect of TMB. **(A)** Example multiplex immunofluorescence from our treatment-naïve independent cohort of immune-classified ovarian cancers showing representative cases of inflamed, excluded, and desert tumors^10^. **(B)** The inflammation score in our independent cohort (see panel A) showed a higher abundance of inflamed tumors in the top tertile and excluded tumors in the bottom tertile. The tertiles are separated with a dashed line. For some patients, multiple samples were sequenced with bulk RNA-seq, and due to intratumor heterogeneity, there are differences across samples from a single patient. Note that in patients with high heterogeneity, the mIF-based classification and bulk RNA-seq may capture different tumor populations and show higher disagreement. **(C)** TMB distribution between MMRP (including subgroups HRP, HRD, and HRD-low) samples in dHpC, dCpH, and other tumor types is compared (t-test; p-values shown). The MMRP samples are shown separately. **(D)** TMB distributions between HRD and HRP samples are compared within each tumor type group (dHpC, dCpH and others). **(E)** T values (as indicated by fill colors) obtained from the multi-linear model for Z-normalized IS using HRD status and TMB as covariates performed on groups of tumor types (dHpC, dCpH and others). *p: 0.05-0.01, 0.01-0.001, 0.001-0.0001, and < 0.0001* are indicated by ‘.’, ‘*’, ‘**’, ‘***’, and *‘****’*.

**Figure S4.**
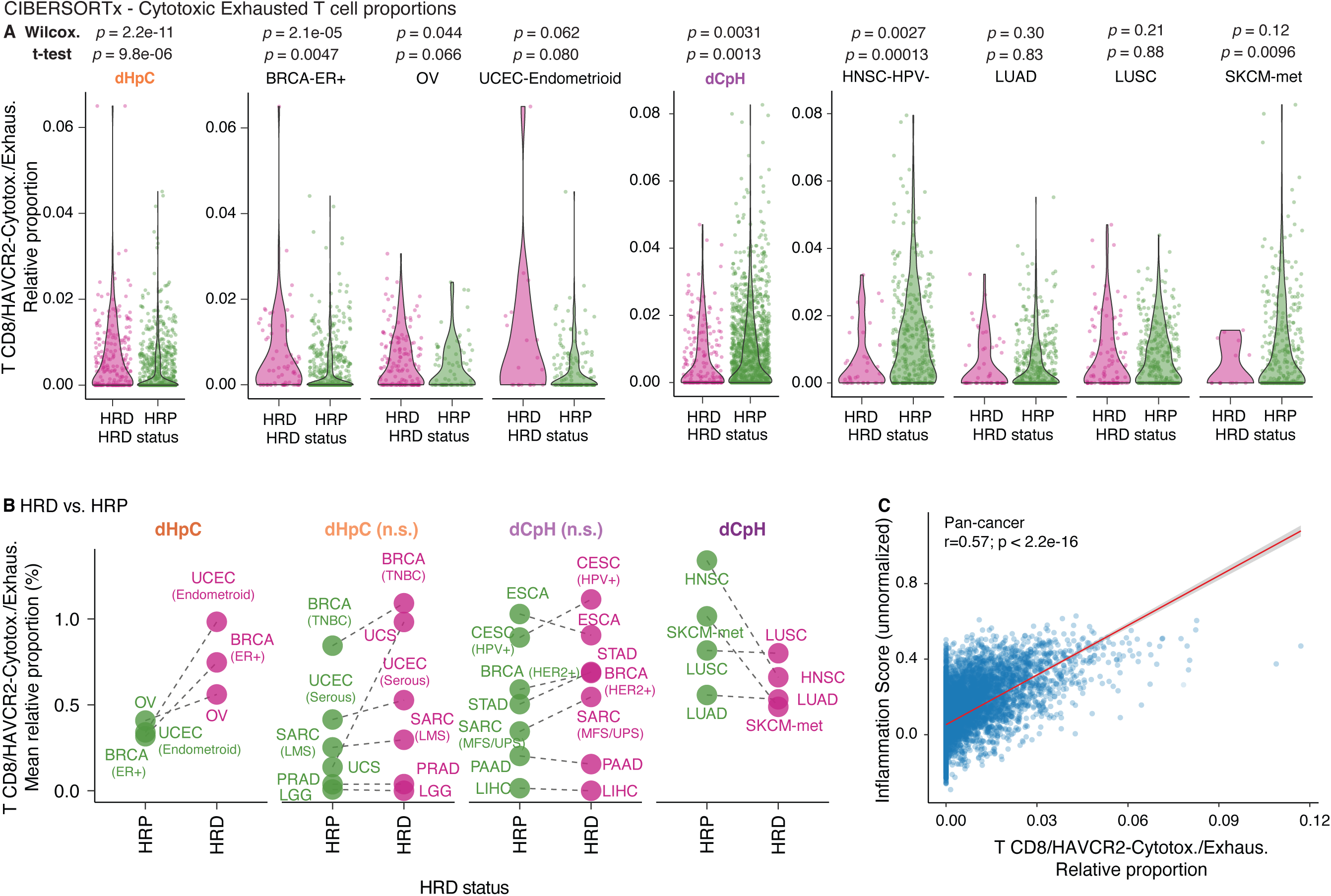
Cytotoxic Exhausted CD8+ T cells across tumor types and correlations with IS. **(A)** The distribution of relative proportions of cytotoxic/exhausted HAVCR2^+^ CD8^+^ T cells as inferred by CIBERSORTx. Proportions in HRD and HRP tumors are compared by the Wilcoxon test and t-test (*p* values are shown on top) for dHpC and dCpH groups combined and individual tumor types in these groups. **(B)** The tumor type-averaged values of the HAVCR2^+^ CD8^+^ T in dHpC, dHpC-n.s., dCpH-n.s., and dCpH. **(C)** Correlation between inflammation score (no tumor type-specific Z-score normalization) and HAVCR2^+^ CD8^+^ T cell relative proportions. Pearson correlation test was performed.

**Figure S5.**
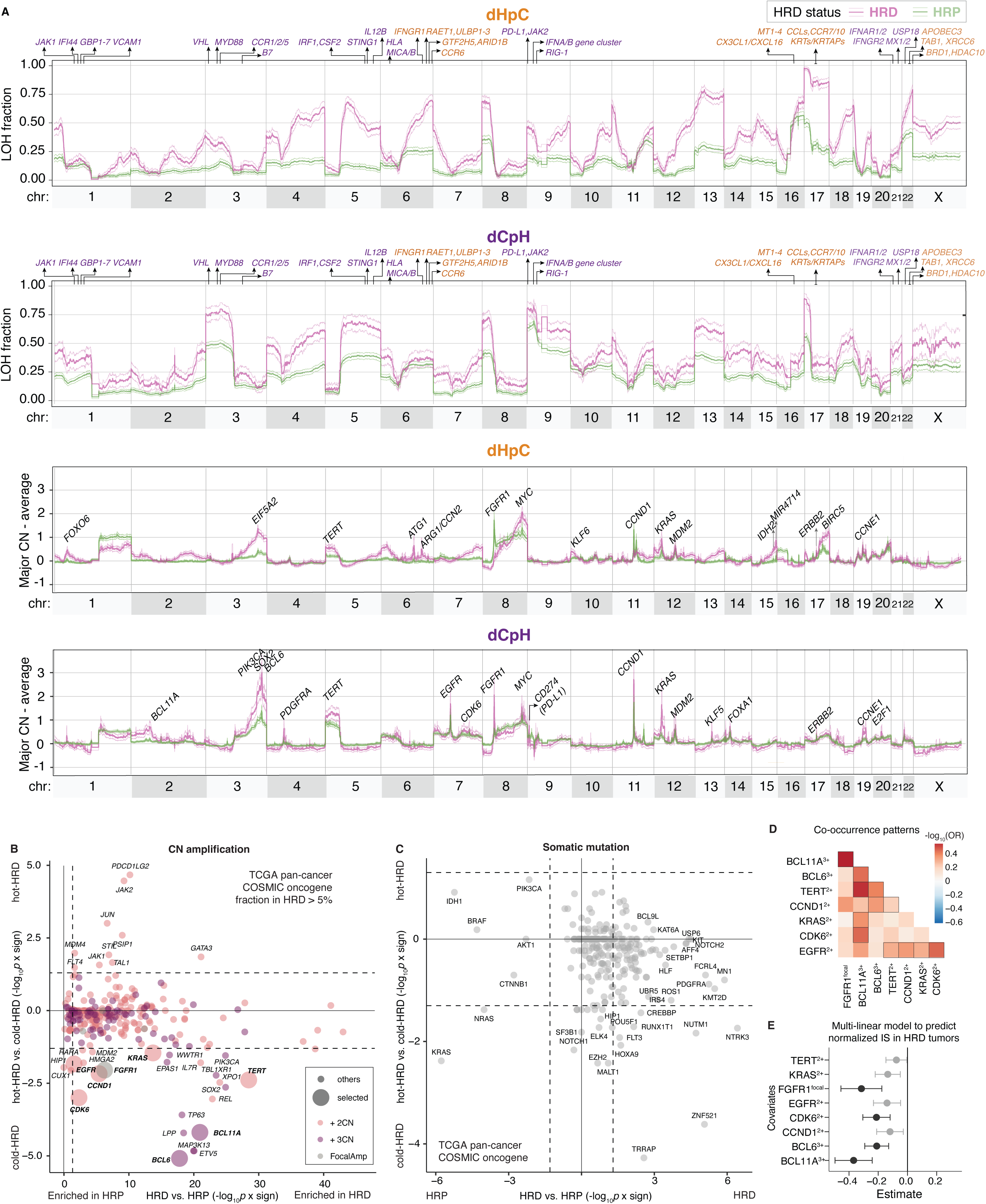
Copy number alteration profiles and their differences between HRD and HRP tumors. **(A)** Same as in Figure 4 but comparing HRD and HRP tumors from top to bottom: (i) Fraction of samples with LOH in dHpC (i) and dCpH (ii), amplification profiles as defined by major CN (after subtracting mean major CN across chromosomes) in dHpC (iii) and dCpH (iv). **(B)** COSMIC oncogenes with amplifications (considering 2+, 3+ for major CN with respect to the baseline and focal amplifications; indicated by marker colors) in more than 5% of pan-cancer HRD samples were identified. The frequencies were compared between (i) hot HRD vs. cold HRD tumors, and (ii) HRD vs. HRP tumors were compared. The figure shows the scatter plot for -log_10_p-values scaled by the sign of Log-Odds Ratio (LOR) from the two comparisons. The genes enriched in HRD and cold HRD tumors were selected, in regions with multiple oncogene amplifications, the one with higher significance in hot vs cold HRD comparison was picked (*e.g*. BCL6, PIK3CA, and SOX2 are all located nearby at chr3 q26.33-q27 loci and often co-amplified). The ones selected are marked with larger marker sizes. **(C)** Same as in panel B but showing oncogenic mutations.

**Figure S6.**
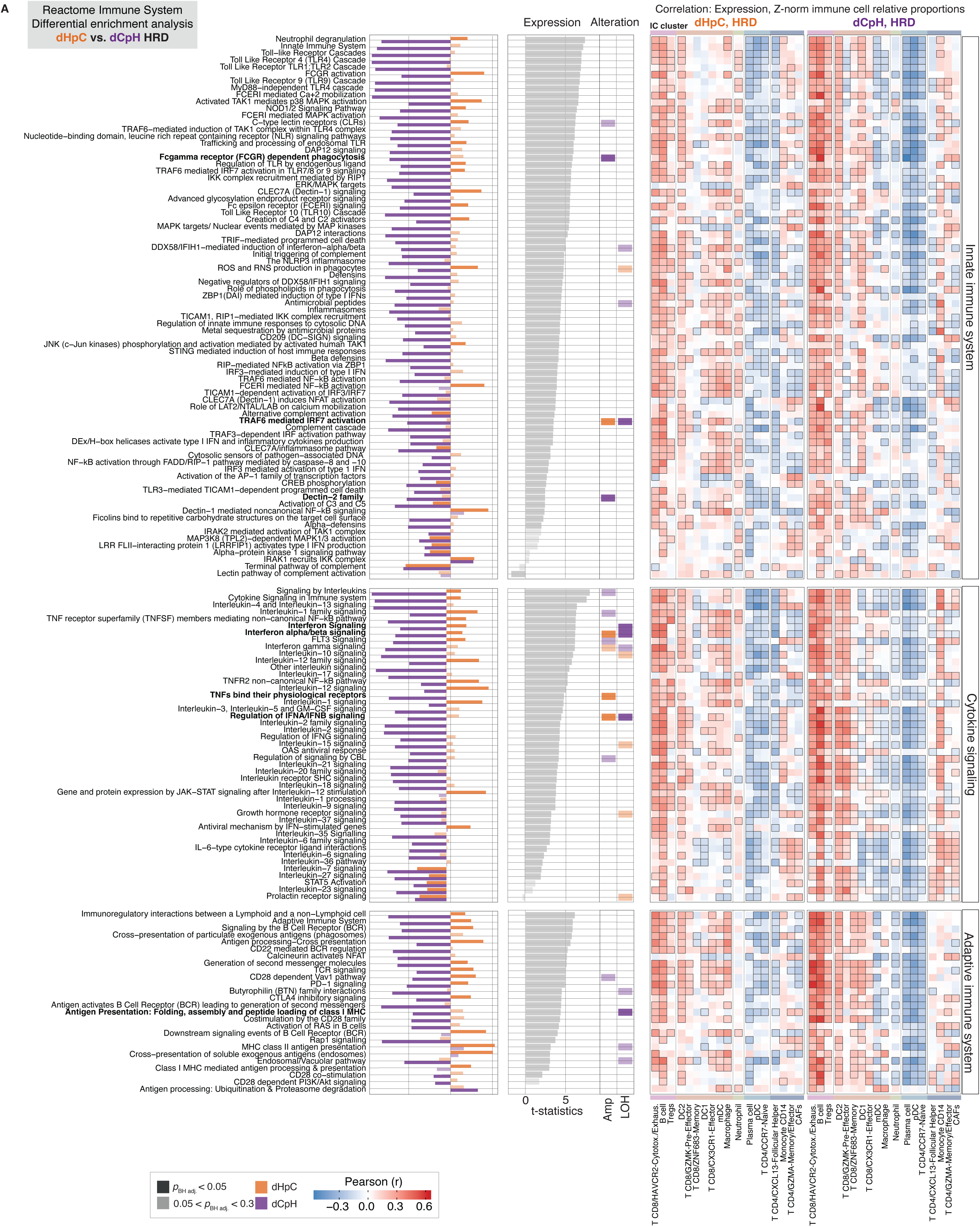
Multi-modal enrichment and correlative analysis for all Reactome immune signaling pathways. Same as Figure 5A-F but for an extended set of pathways.

**Figure S7.**
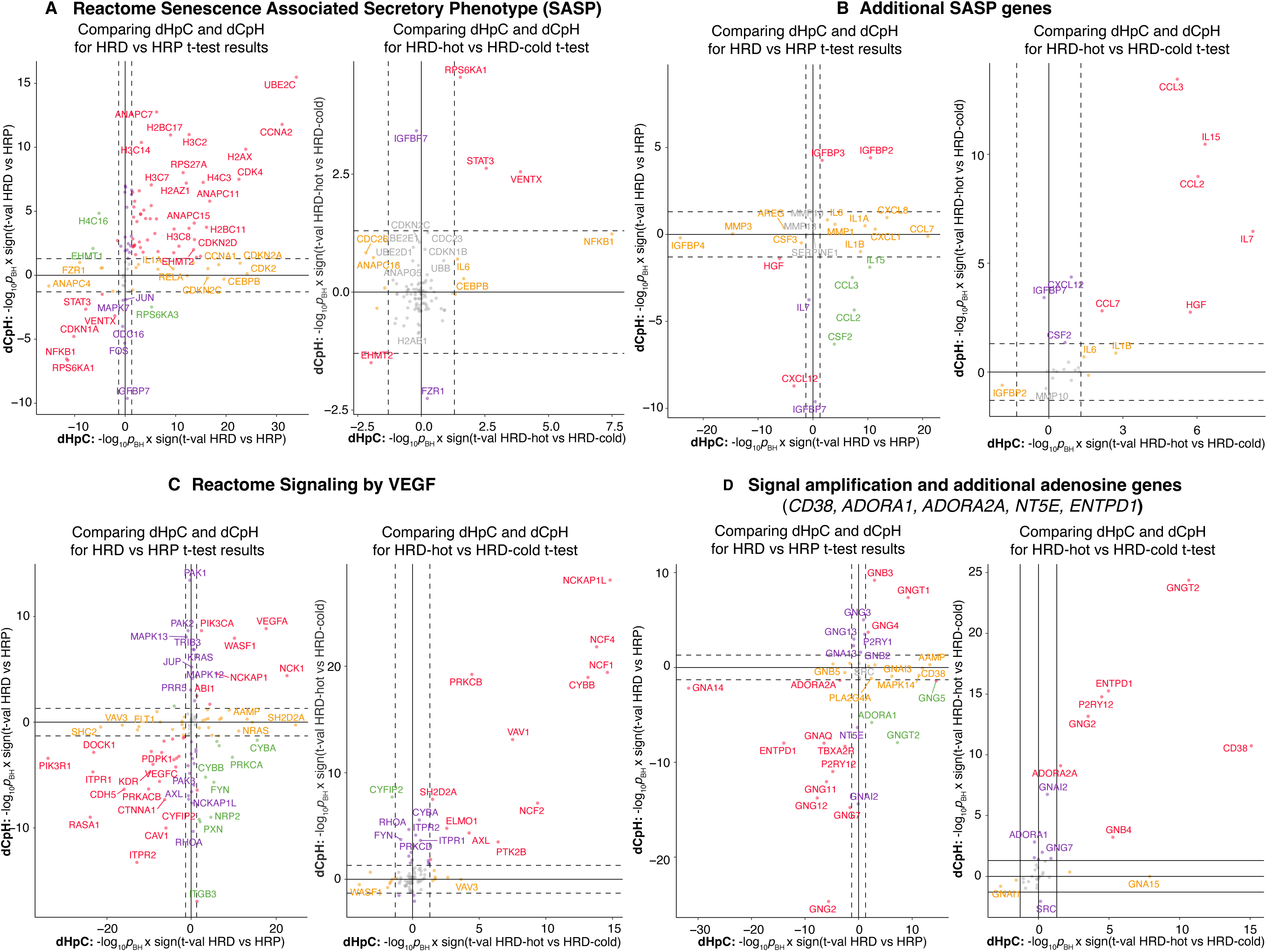
Association between gene expression of selected pathways and HRD tumor archetypes. **(A**-**D)** For each panel, gene expression for genes in a given pathway was compared between HRD and HRP samples for dHpC and dCpH (left subpanels), and for HRD-hot vs HRD-cold samples in dHpC and dCpH using t-test. The x-axis shows results for dHpC in the form of -log_10_p scaled by the sign of t value. Red indicates significant in both tumor type groups, orange in dHpC and purple in dCpH only, green indicates opposite modifications in the two tumor type groups. Results are shown in order for Reactome Senescence Associated Secretory Phenotype (SASP) pathway (R-HSA-2559582) **(A)**, additional SASP genes manually curated and not included in the reactome pathway **(B),** Reactome Signaling by VEGF (R-HSA-194138) **(C)** and signal amplification (R-HSA-392518) pathways together with additional adenosine genes (*CD38*, *ADORA1*, *ADORA2A*, *NT5E*, and *ENTPD1*) **(D)**.

